# Epidermal barrier dysregulation in atopic skin predisposes for excessive growth of the allergy-associated yeast *Malassezia*

**DOI:** 10.1101/2023.09.11.557156

**Authors:** Fiorella Ruchti, Pascale Zwicky, Burkhard Becher, Sandrine Dubrac, Salomé LeibundGut-Landmann

## Abstract

The skin barrier is vital for protection against environmental threats including insults caused by skin-resident microbes. Dysregulation of the barrier is a hallmark of atopic dermatitis (AD) and ichthyosis, with variable consequences for host immune control of colonizing commensals and opportunistic pathogens. While sensitisation to *Malassezia*, the most abundant commensal fungus of the skin, is common in AD, its relevance for pathogenesis remains unclear. Here we show that in barrier-disrupted skin, *Malassezia* acquires enhanced fitness. This is not a consequence of the dysregulated allergic immune status characteristic for AD but is rather explained by structural and metabolic changes in the cutaneous niche that provide increased accessibility and a favourable lipid profile, to which the lipid-dependent yeast adapts for enhanced nutrient assimilation. These findings reveal fundamental insights into the role of the mycobiota in the pathogenesis of common skin barrier disorders.

## INTRODUCTION

The microbiota is an integral part of the skin ecosystem that enhances the barrier function of this essential organ ensuring protection of the host from dehydration and entry of harmful substances including toxins and pathogens.^1,2^ The essential role of commensal microbes in skin homeostasis is illustrated by the dysregulation of physical, chemical, and immunological properties of the cutaneous barrier under germ-free conditions.^3,4^ Disease of the skin are commonly accompanied by dysbiosis of the cutaneous microbiota, especially in pathologies bearing barrier defect such as atopic dermatitis (AD) and ichthyosis.^5,6^ The functional relationship between the microbiota and pathogenesis is however not well defined.

AD is a very prevalent chronic relapsing skin disorder with with a high disease burden and an increasing incidence, in both, humans and dogs.^7,8^ Due to its highly pruritic and eczematous manifestation, AD can have a profound impact on patients’ quality of life. Furthermore, AD frequently progresses to other allergic conditions including food allergy, allergic rhinitis and/or asthma, also referred to as the atopic march.^9^ The complex etiology of AD involves genetic and environmental factors that are associated with barrier defects, immune dysregulation and infection. It is believed that impaired barrier allows penetration of microbial and environmental antigens to deeper layers of the epidermis. This promotes allergen sensitization and inflammation, which perpetuate damage to the epithelial barrier integrity via an inflammatory amplifying loop.^10^ As such, *Staphylococci* accumulate in AD lesions and produce toxins that exacerbate epidermal inflammation and barrier disruption.^11,12^ Another skin microbe that has been tied to AD is *Malassezia*,^13^ although its causal relationship to the disease is less well established. *Malassezia* is the most abundant fungus in normal human skin and in many warm-blooded animals.^14^ Of the 20 species described so far, *M. restricta* and *M. globosa* dominate human skin, followed by *M. sympodialis* and *M. furfur*, while *M. pachydermatis* is the most abundant species on canine skin and a common agent of otitis externa in this host species.^15–17^ Although generally viewed as a commensal, *Malassezia* is associated with diverse pathological skin conditions, including AD.^13,18^ More recently, *Malassezia* was even connected to extracutaneous pathologies including IBD^19^ and cancer.^20,21^ In AD, about half of all adult patients, including preferentially those with head- and neck dermatitis, are sensitized to *Malassezia*.^22,23^ In veterinary dermatology, *M. pachydermatis* sensitization is observed in more than one third of dogs with canine AD.^24^ Several *Malassezia*-derived allergens have been characterized. While *Malassezia*-specific IgE antibodies and Th2 cells have been proposed to cross-react with conserved human antigens and to mediate self-reactivity, their role in AD pathogenesis remains unclear.^25,26^ Likewise, the implications of *Malassezia* dysbiosis that has been reported in AD lesional skin is unknown.^27,28^

*Malassezia* is highly adapted for life on the mammalian skin. Given its dependence on exogenous lipids due to the lack of fatty acid synthase, *Malassezia* finds favorable conditions in lipid-rich areas of the skin such as sebaceous hair follicles.^29^ By metabolizing skin lipids for its own benefit, the fungus contributes to the lipid balance of the cutaneous niche. In turn, *Malassezia* is tightly controlled by the cutaneous immune system, which prevents fungal overgrowth. Hereby, the IL-17 pathway is key for ensuring homeostatic colonization.^30^

How *Malassezia* commensalism is affected under conditions of skin inflammation and barrier disruption is however not well defined. To address this, we employed four different experimental models of AD and cutaneous barrier stress and assessed how changes in the immune status, barrier integrity, and lipid availability affect *Malassezia* colonization. We provide evidence that *Malassezia* acquires enhanced fitness in the dysfunctional skin barrier. More specifically, we show that unrestrained *Malassezia* growth in the AD-like skin is independent of the type 2-polarized immune environment, but instead is linked to barrier disruption and alterations in the lipid metabolism in the dysregulated skin barrier to which the fungus adapts for enhanced lipid assimilation.

## RESULTS

### *Malassezia* grows excessively in the atopic skin environment

To study the interaction of *Malassezia* with the host under AD-like conditions we adopted the widely used MC903 model of AD in C57BL/6 mice.^31,32^ *Malassezia pachydermatis* or olive oil (vehicle) was applied to the dorsal ear skin^33^ after induction of an AD-like immune phenotype by repeated application of the vitamin D analogue calcipotriol (MC903) or ethanol (EtOH; solvent control) for ten days (Fig. 1A). MC903 induced an AD-like phenotype characterized by an increase in ear thickness (Fig. 1A), epidermal hyperplasia (Fig. 1B-C), elevated skin *Tslp* transcripts (Fig. 1D) and serum IgE titers (Fig. 1E). Epicutaneous application of *M. pachydermatis* slightly worsened epidermal hyperplasia (Fig. 1B) but did not further increase the overall ear thickness by day 4 and day 7 post-infection (Fig. 1A, C). At these time points *Tslp* transcript levels were reduced in response to *Malassezia* association (Fig. 1D). Of note, maintenance of the AD-like phenotype required continued application of MC903 (Fig. 1A). To minimize interference of the ethanol solvent with fungal viability, the continuous treatment after infection was applied to the ventral side of the ear only (Fig. 1A, Fig. S1A-C). The unilateral treatment with MC903 led to a robust expression of the vitamin D receptor target gene *Cyp24a1* on both dorsal and ventral ear sides and did not affect fungal loads or inflammatory infiltrates (Fig. S1A-C).

**Fig 1.**
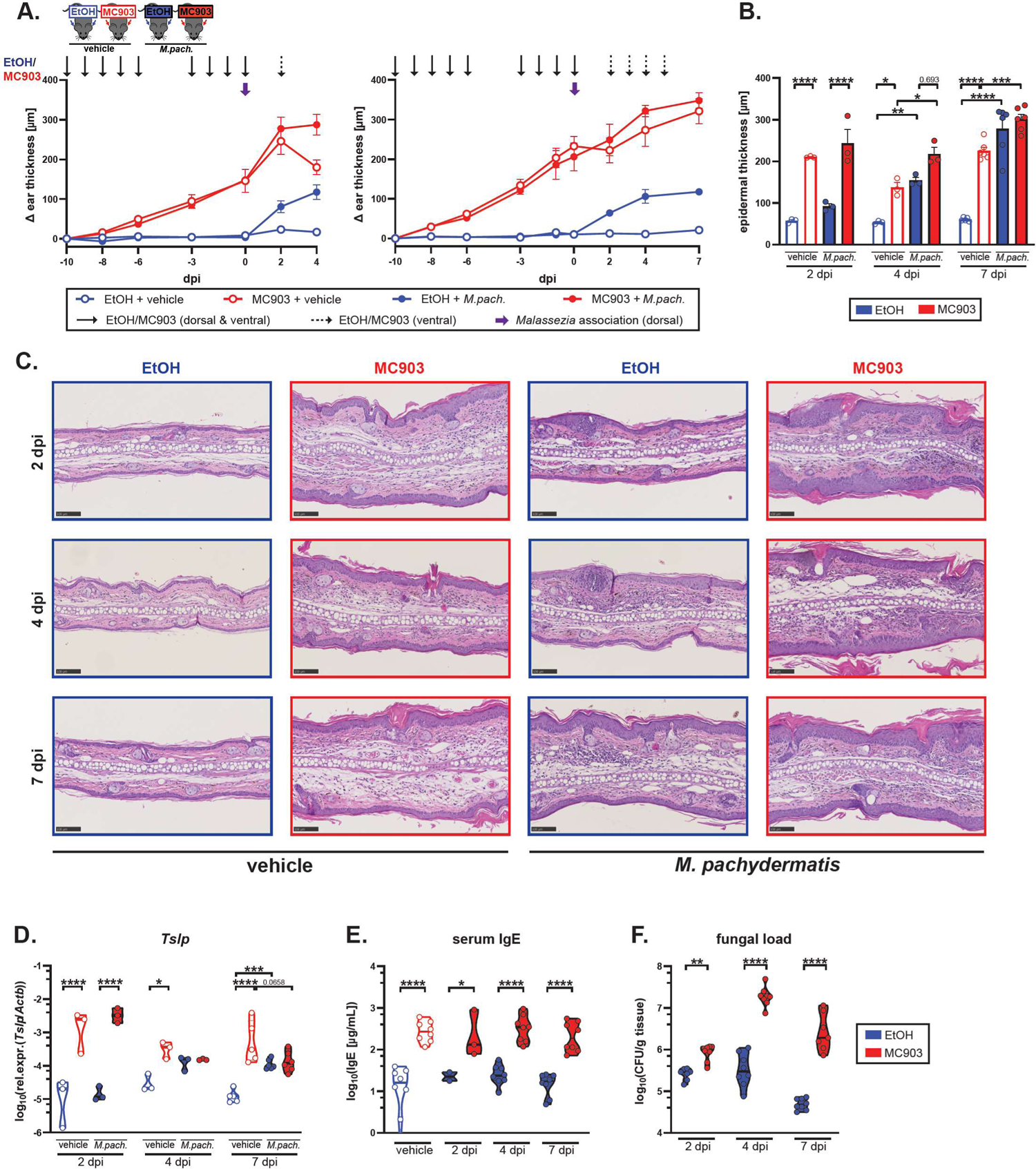
*Malassezia* overgrowth in the atopic skin environment. **A.-F.** The ear skin of WT C57BL/6 mice was repeatedly treated with MC903 (red) or ethanol as a control (EtOH, blue) for 10 days, and then associated dorsally with *Malassezia pachydermatis* (*M.pach.*, closed symbols) or olive oil (vehicle, open symbols) for 2, 4 or 7 days (dpi). **A.** Ear thickness kinetics. The experimental schedule is indicated at the top of the panel with long arrows representing MC903 or EtOH administration and wide arrows representing *M. pach.* association. **B.-C.** epidermal thickness (B) and representative H&E-stained ear sections (C) at the indicated time points. **D.-F.** *Tslp* transcript levels in the skin (D), serum IgE levels (E), and skin fungal load (CFU) (F). Data are pooled from two (A-D), three (F), or four (E) independent experiments with 3-4 mice per group each. Each datapoint represents one mouse (B, D, E, F) or indicates the mean +/- SEM (A) at the indicated time points. Horizontal bars in D-F indicate the mean of each group. Statistical significance was determined using two-way ANOVA. *p<0.05, **p<0.01, ***p<0.001, ****p<0.0001. **See also Figure S1.**

We then assessed the fate of *Malassezia* in the AD-like inflammation and found that the altered skin environment strongly interfered with fungal control. We observed pronounced fungal overgrowth in the MC903-treated skin compared to *Malassezia*-colonized control skin as early as 2 dpi with a further increase over the course of 7 days, while control mice steadily controlled the fungus (Fig. 1F). The elevated fungal burden in MC903-pretreated skin was not limited to a single species of *Malassezia* but was also seen after infection with *M. sympodialis* and *M. furfur* (Fig. S1D). Moreover, the same was observed in mice on the Balb/c background, which are widely used in allergy research due to their inclination to type 2 immune polarization (Fig. S1E-G). Taken together, these data show that while *Malassezia* does not aggravate atopy in MC903-treated mice, it strongly overgrows under these conditions.

### Selected *Malassezia*-induced immune populations are modulated in the MC903-treated skin

Following the marked fungal overgrowth in the AD-like skin, we hypothesized that the host response against *Malassezia* may be compromised. We therefore investigated potential alterations in the immune status of the MC903-treated skin. By high dimensional spectral flow cytometry, we characterized and quantified changes in the immune cell composition of the skin in response to topical MC903. Populations were defined by unsupervised analysis using tSNE clustering and quantification of individual cell types by classical gating (Fig. 2A, B, Fig. S2A). As characteristic for AD skin lesions and previously described for the MC903-AD mouse model,^31^ we observed an accumulation of diverse inflammatory immune cells (Fig. 2, Fig. S2) as also evidenced in H&E-stained histology sections (Fig. 1C).

**Fig 2.**
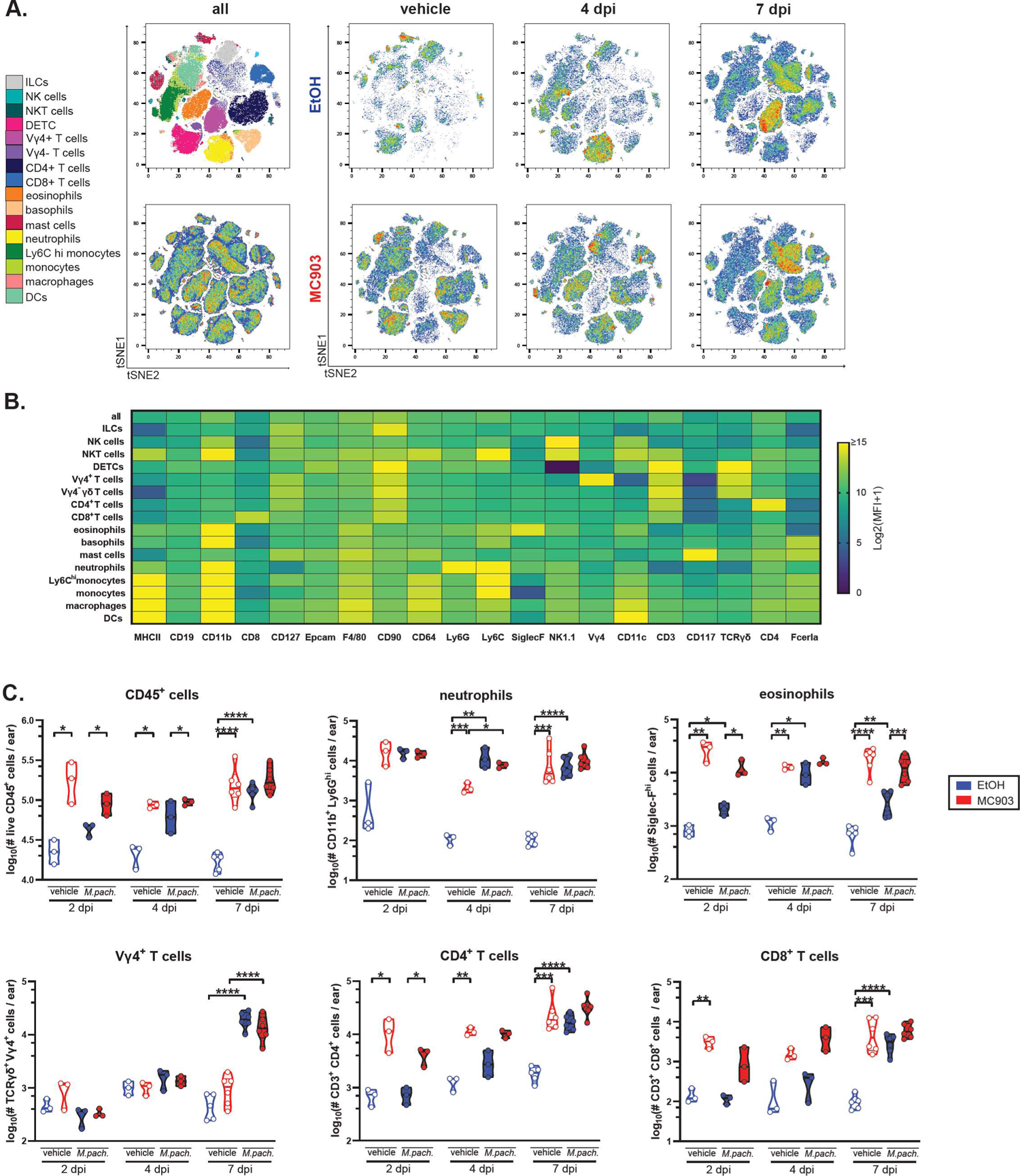
MC903-induced skin inflammation is characterized by infiltration of diverse immune cell subsets. Immunophenotyping of CD45^+^ cells in the ear skin of WT C57BL/6 mice treated with MC903 or EtOH and associated with *M. pachydermatis* or vehicle for 2, 4 or 7 days (dpi) as in Fig. 1A. **A.** tSNE plots of all samples combined (all) or of three concatenated samples per indicated treatment group in naïve (vehicle) and *M. pachydermatis*-exposed mice at 4 or 7 dpi. Color overlays correspond to manually gated cell populations shown on the left. **B.** Heat map indicating the relative expression of markers per identified cell type. **C.** Quantification of the identified cell populations at the indicated time points after fungal association or vehicle application to MC903-treated or EtOH-treated control skin. Each symbol represents one mouse. The mean of each group is indicated. Data are pooled from two independent experiments with three mice per group each. Statistical significance was determined using two-way ANOVA. *p<0.05, **p<0.01, ***p<0.001, ****p<0.0001. **See also Figure S2.**

*Malassezia* colonization of the skin resulted in an accumulation of hematopoietic cell subsets with a gradual increase over time, independently of MC903. At early time points they were dominated by myeloid cells with an early sharp increase in neutrophils at 2 dpi followed by inflammatory monocytes, basophils, and eosinophils accumulating at 4 dpi (Fig. 2A, C, Fig. S2B). T cell numbers increased only by 7 dpi including cutaneous CD4^+^ and CD8^+^ subsets and, most prominently and selectively, Vγ4^+^ γδ T cells (Fig. 2A, C).

When comparing *Malassezia*-induced immune cell subsets in the MC903-treated skin to those in control-treated skin we found several known and suspected anti-fungal effector cell populations to be increased under AD-like conditions, including CD4^+^ and CD8^+^ T cells, eosinophils, and basophils (Fig. 2A, C, Fig. S2B). Of note, no *Malassezia*-induced cell populations were diminished in the skin of MC903-treated compared to control-treated mice. Together, these changes in immune cell population dynamics are unlikely to provide an explanation for the observed *Malassezia* overgrowth in AD-like skin, which leaves putative functional changes in effector cells as possible cause of uncontrolled fungal expansion.

### *Malassezia* dysbiosis in the MC903-treated skin is independent of type 2 immunity

Given the pronounced induction of TSLP and other hallmarks of type 2 immunity in MC903-treated skin, we scrutinized this immune pathway as a potential cause of *Malassezia* overgrowth. TSLP is the main driver of the type 2 response in the MC903 model of AD. ^34,35^ Consequently, the number of IL-4^+^ T cells in the draining lymph nodes of MC903 treated mice was increased upon MC903 treatment (Fig. 3A). Interestingly, the *Malassezia*-responsive subset of CD4^+^ T cells elicited in MC903-treated mice did not produce Th2 cytokines when re-stimulated in an antigen-specific manner, despite the pronounced type 2 polarizing environment, but instead gave rise to Th17 effector cells as observed in the solvent control group (Fig. S3A-D).

**Fig 3.**
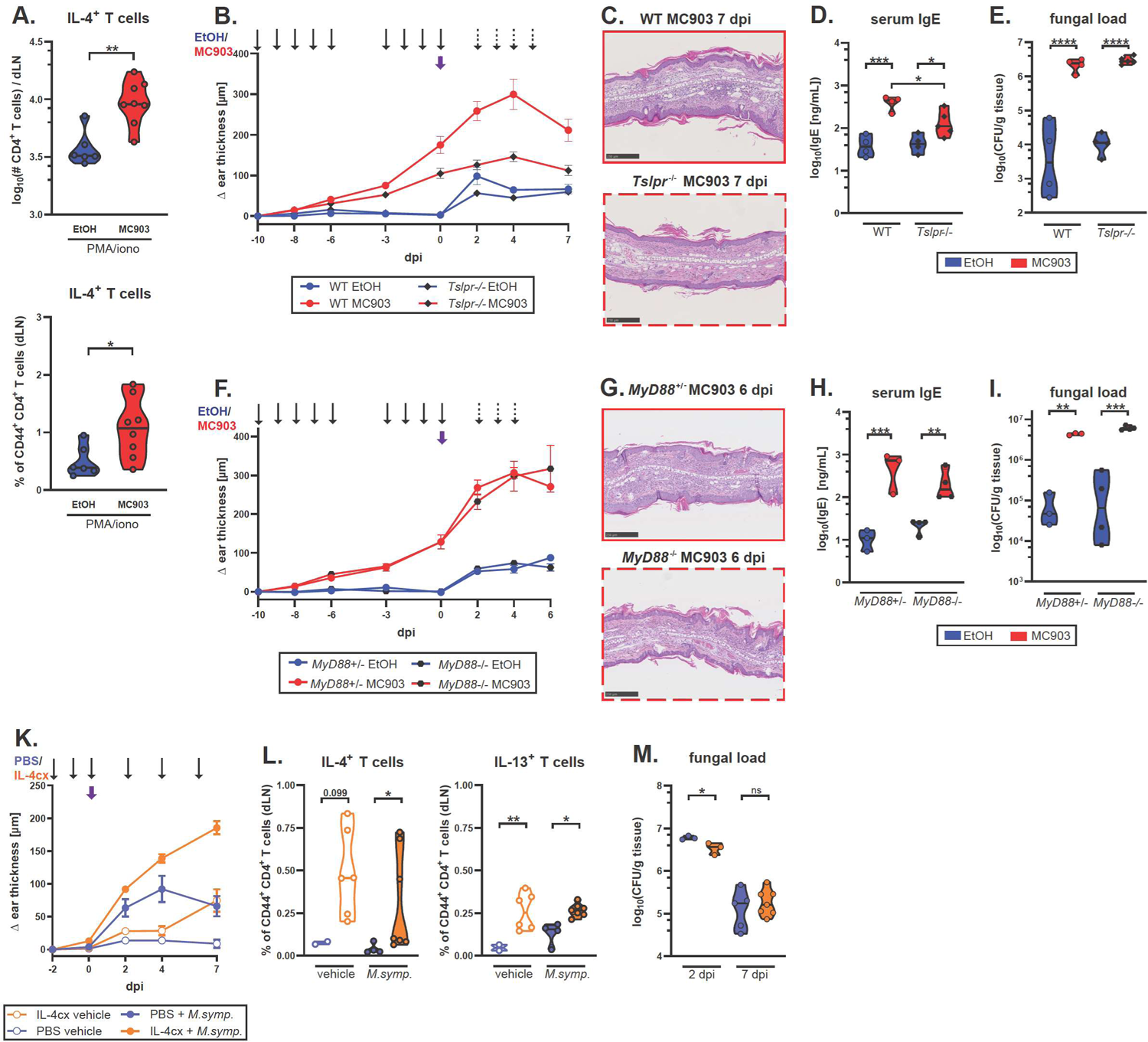
Fungal overgrowth in the MC903-treated skin is independent of type 2 immunity. A. The ear skin of WT C57BL/6 mice was treated with MC903 or EtOH and associated with *M. pachyermatis* as in Fig. 1A. IL-4 production by dLN CD4^+^ T cells at 7 dpi was quantified after *ex vivo* re-stimulation with PMA and ionomycin. Data are pooled from two independent experiments with 3-4 mice per group each with each symbol representing one mouse, the mean is indicated. **B.-I.** The ear skin of *Tslpr*^-/-^ and WT control mice (B-E) or *MyD88*^-/-^ and *MyD88*^+/-^ control mice (F-I) was treated with MC903 or EtOH and associated with *M. pachydermatis* (*M.pach.*) as in Fig. 1A for seven or six days, respectively. The experimental schedule is indicated at the top of (B) and (F). Ear thickness kinetics (B, F), representative H&E-stained skin sections (C, G), serum IgE levels (D, H), and skin fungal load (E, I). Data in B-E are from one representative of two independent experiments. Data in F-I are from one experiment. In B and F, each symbol is the mean +/- SEM (B) or mean +/- SD (F) of each group at the indicated time points. In D, E, H, and I, each symbol represents one mouse, the mean of each group is indicated. **K.-M.** WT C57BL/6 mice were injected intraperitoneally with mIL-4 complexed with anti-mIL-4 mAb (IL-4cx; orange) or PBS (blue) and the ear skin was associated dorsally with *M. sympodialis* (*M.symp.*, closed symbols) or vehicle (open symbols). The experimental schedule is indicated at the top of (K) with thin arrows representing IL-4cx or PBS administration and a wide arrow representing *M.symp.* skin association. Ear thickness kinetics (K), quantification of IL-4^+^ and IL-13^+^ CD4^+^ T cells in skin draining lymph nodes after *ex vivo* PMA and ionomycin re-stimulation at 7 dpi (L) and skin fungal load at 2 and 7 dpi (M). Data are pooled from two independent experiments (one experiment only for the 2 dpi time point in M), with 2-4 mice per group each. In K, each symbol is the mean +/- SEM per group and time point. In L and M, each symbol represents one mouse, the mean of each group is indicated. Statistical significance was determined using Student’s unpaired *t*-test (A), one-way ANOVA (L, M) or two-way ANOVA (D, E, H, I). *p<0.05, **p<0.01, ***p<0.001, ****p<0.0001. **See also Figure S3.**

We then made use of *Tslpr*^-/-^ mice^36^ to assess the impact TSLP and downstream effects on fungal control in response to MC903. Absence of TSLP signaling resulted in a decrease of overall ear inflammation after MC903 treatment as reflected by the diminished ear thickness (Fig. 3B), the reduced number of inflammatory cells infiltrating the skin (Fig. 3C, Fig. S3E) and the lower serum IgE levels (Fig. 3D). Importantly, these changes had no effect on fungal control when comparing MC903-pre-treated *Tslpr*^-/-^ and wild-type (WT) mice (Fig. 3E). These results were replicated in *MyD88*^-/-^ mice lacking IL-1 family cytokine signaling including IL-33 and IL-18 signaling, which were reported to be involved in driving the MC903-induced type 2 skin inflammation.^37,38^ In comparison to *Tslpr*^-/-^ mice, MyD88-deficiency had a limited effect on the AD-like skin phenotype as indicated by the preserved marked ear inflammation (Fig. 3F-G) and elevated serum IgE levels (Fig. 3H). Fungal overgrowth in response to MC903 was unchanged in comparison to MyD88-sufficient control animals (Fig. 3I).

To further investigate the impact of type 2 immunity on fungal control in a MC903- and TLSP-independent context, we repeatedly injected WT mice with recombinant IL-4 to increase systemic IL-4 concentrations before and during skin colonization with *Malassezia* (Fig. 3K). IL-4 was administered in the form of IL-4-anti-IL-4 monoclonal antibody complexes (IL-4cx), which prolongs the cytokine’s *in vivo* biological half-life.^39^ While IL-4cx injection resulted in an increase in epidermal and overall ear thickness (Fig. S3F) and percentages of IL-4- and IL-13-producing CD4^+^ T cells (Fig. 3L), the fungal load remained unaffected at both 2 and 7 dpi in comparison to the PBS-injected control group (Fig. 3M), indicating that systemic increase in type 2 cytokines did not interfere with fungal control and is therefore unlikely the cause of fungal overgrowth under AD-like conditions. Together, these results indicate that local or systemic type 2-biased immunity is not per se responsible for impaired fungal control in the AD-like skin.

### Decreased type 17 antifungal immunity alone does not explain *Malassezia* overgrowth in MC903-treated skin

IL-17 immunity is vital for antifungal defense in barrier tissues including cutaneous protection against *Malassezia* in experimentally colonized mice.^30^ Therefore, we aimed at assessing the role of IL-17 in fungal control in MC903-treated skin. We found reduced transcript levels of *Il17a* and *Il17f* at 4 and 7 days after *M. pachydermatis* association (Fig. 4A). Cutaneous IL-17A producing cells were reduced in numbers as well as in the extent of cytokine production per cell (Fig. 4B-C, Fig. S4A), whereas IFN-y-production was not affected (Fig. S4B). Among the IL-17A-producing T cell subsets, Vy4^+^ yd T cells showed the highest IL-17A production in response to *M. pachydermatis* and, in turn, the most pronounced reduction in the AD-like skin compared to control skin (Fig. 4C). This drop in IL-17 production was further apparent in *Malassezia*-responsive IL-17A^+^ Vy4^+^ yd T cells in the ear draining lymph nodes (Fig. S4C).

**Fig 4.**
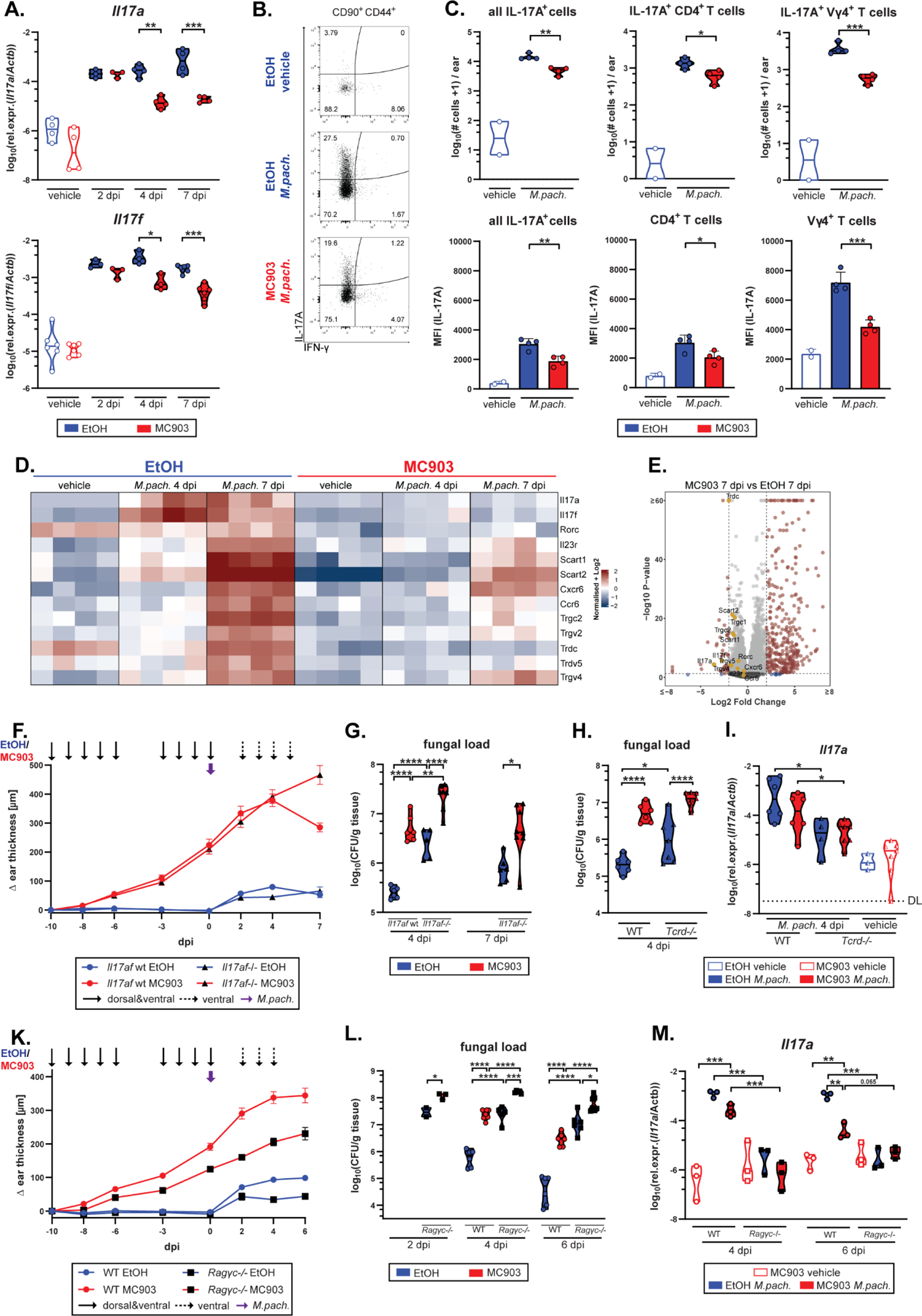
Decreased type 17 immunity in the MC903-exposed skin alone is not responsible for fungal overgrowth. **A.-C.** The ear skin of C57BL/6 mice was treated with MC903 or EtOH and associated with *M. pachydermatis* or vehicle as in Fig. 1A and IL-17 expression was analyzed in the skin. Quantification of *Il17a* and *Il17f* transcript levels at 2, 4, and 7 dpi (A), representative flow cytometry plots showing IL-17A and IFN-γ expression by CD90^+^ CD44^+^ T cells from two (vehicle) or four (*M.pach.*) concatenated samples (B), and flow cytometry-based quantification of IL-17-producing cell subsets (top) and IL-17A median fluorescence intensities (MFI; bottom) as indicated, at 7 dpi. Each symbol in A and C represents one mouse, the mean of each group is indicated. Data are from one of two (B-C) or from two (A) independent experiments with 2-4 mice per group each. **D.-E.** The ear skin of C57BL/6 mice was treated with MC903 or EtOH and associated with *M.pach.* or vehicle as in Fig. 1A and bulk ear skin RNA was sequenced at 4 and 7 dpi and from the vehicle group. Heat map of selected IL-17- and γδ T cell-associated genes displayed as log2 fold change (D). Each column is one mouse. Volcano plot (E) with differentially expressed genes (DEGs) in MC903-treated vs. EtOH control skin at 7 dpi. Significant DEGs (log2 fold change >2; p<0.05) are displayed in red, IL-17- and γδ T cell-associated genes are highlighted in gold. **F.-M.** The ear skin of *Il17af*^-/-^ and *Il17af*^wt^ mice (F-G), *Tcrd^-/-^* and WT control mice (H-I) or *Rag2*^-/-^ *Il2rg*^-/-^ (*Ragγc*-/-) and WT control mice (K-M) was treated with MC903 or EtOH and associated with *M. pachydermatis* or vehicle as in Fig. 1A. The experimental schedule is indicated at the top of (F) and (K). Ear thickness kinetics (F, K), skin fungal load (G, H, L), and *Il17a* transcript levels at the indicated time points. DL, detection limit. In F and K, each symbol is the mean +/- SEM per group and time point. In G-I, L, and M, each symbol corresponds to one mouse, the mean of each group is indicated. Data are pooled from two independent experiments with 3-4 mice per group each. Data in L are from one representative of two independent experiments, with the exception of the 2 dpi time point, which is from a single experiment with 2-3 mice per group. Statistical significance was determined using one-way ANOVA (C) or two-way ANOVA (A, G-I, L-M). *p<0.05, **p<0.01, ***p<0.001, ****p<0.0001. **See also Figure S4.**

RNA sequencing of bulk MC903-treated skin associated with *M. pachydermatis* and of corresponding controls revealed the differential expression of many genes in AD-like compared to ethanol skin. More specifically, 341 genes were upregulated at 7 dpi in the MC903 condition while only 44 were downregulated (FDR<0.05, Log2 fold change>2) (Fig. 4D-E). Among the downregulated genes, a strong reduction in the expression of fungus-induced cytokines namely *Il17a* and *Il17f* was observed. In addition, genes associated with IL-17-producing γδ T cells were repressed under AD-like skin conditions (Fig. 4D-E). These included the genes encoding the T cell receptor gamma (*Trgc2*, *Trgv2*, *Trgv4*, *Trgv*5) and delta (*Tcrdc*) chains as well as γδ T cell-associated scavenger receptors (*Scart1*, *Scart2*), the transcription factor Rorγt (*Rorc*), the IL-23 receptor (*Il23r*) and chemokine receptors assigned to IL-17A-producing T cells (*Ccr6*, *Cxcr6*).^40^ In line with previous reports on vitamin D and its analogue calcipotriol (MC903) transcriptionally repressing the IL-17A locus and inhibiting accumulation of IL-17^+^ T cells in psoriasis and autoimmune encephalitis,^41–45^ we found that the decrease in *Il17a* transcript levels in the MC903-treated skin was independent of TSLPR or MyD88-signaling (Fig. S4D-E).

Next, we investigated whether MC903-dependent IL-17 repression was responsible for the observed fungal outgrowth in the AD-like skin (Fig. 4F). To this end, we assessed the fungal load in the skin of MC903-treated *Il17af*^-/-^ and littermate control mice. In control skin, *Il17af* deletion resulted in an increased fungal burden (Fig. 4G), as we showed previously^30^. The requirement for IL-17A/F in fungal control was also evident in the MC903-treated skin (Fig. 4G). Importantly, fungal burden was increased in MC903-compared to ethanol control-treated skin, regardless of the presence or absence of IL-17A/F. Still, the fold difference in CFU was reduced in *Il17af*^-/-^ mice (9-fold compared to 23-fold in WT mice), supporting a partial contribution of IL-17 repression to fungal overgrowth in the MC903-AD model (Fig 4G). These results were replicated in *Tcrd*^-/-^ mice, lacking γδ T cell-derived IL-17 (Fig. 4H-I), and in *Rag2*^-/-^ *Il2rg*^-/-^ (*Ragγc*^-/-^) mice, devoid of all lymphocyte and innate lymphoid cell (ILC) subsets (Fig. 4K-M). Fungal overgrowth in *Ragγc*^-/-^ mice was massive, while *Il17a* levels were generally very low in these mice, irrespective of treatment (Fig. 4K-M). This confirms that neither IL-17-nor lymphocyte-mediated effects alone are responsible for the observed fungal overgrowth in the AD-like skin. Interestingly and in line with the notion that fungal overgrowth is largely IL-17-independent in the MC903-treated skin, we found reduced *Il17a* levels in IL-4cx-treated mice while fungal clearance was unaffected (Fig. 3K, Fig. S4F). Taken together, our results show that MC903-mediated repression of IL-17 can only partially explain *Malassezia* overgrowth in the atopic skin.

### Physical barrier impairment in AD-like skin favors *Malassezia* overgrowth

We next considered epidermal barrier impairment, a main feature of AD,^46^ as an underlying cause of *Malassezia* dysbiosis in MC903-induced AD model. Interestingly, MC903 induced AD-like symptoms promote skin fungal overgrowth, regardless of alterations in the IL-1-(*Myd88*^-/-^), type 2 (*Tslp^-/-^*) or type 17 (*Il17af*^-/-^; *Tcrd*^-/-^) immune pathway or in mice devoid of the entire lymphoid compartment (*Ragγc*^-/-^) (Fig. 3E,I and Fig. 4G, H, L). Thus, our results reveal little effects of the skin immune system on *Malassezia* overgrowth. The MC903 mouse model of AD is characterized by physical epidermal barrier impairment as showed by increased TEWL and epidermal hyperplasia.^47^ This feature was preserved upon *Malassezia* association and conserved in all the immune deficient mouse lines manifesting fungal overgrowth upon MC903-treatment (Fig. 5A). Consistent with these histopathological changes, keratinocytes were expanded, most prominently the subset of hair follicle keratinocytes, and displayed an activated phenotype (Fig. 5B, Fig. S5A-C). Of note, the stratum corneum including invaginating hyperkeratotic hair follicles are the main niche of *Malassezia* colonization in the skin (Fig. 5C). This suggests that alterations in keratinocytes and especially the epidermal barrier function might underly *Malassezia* overgrowth upon MC903-induced AD.

**Fig 5.**
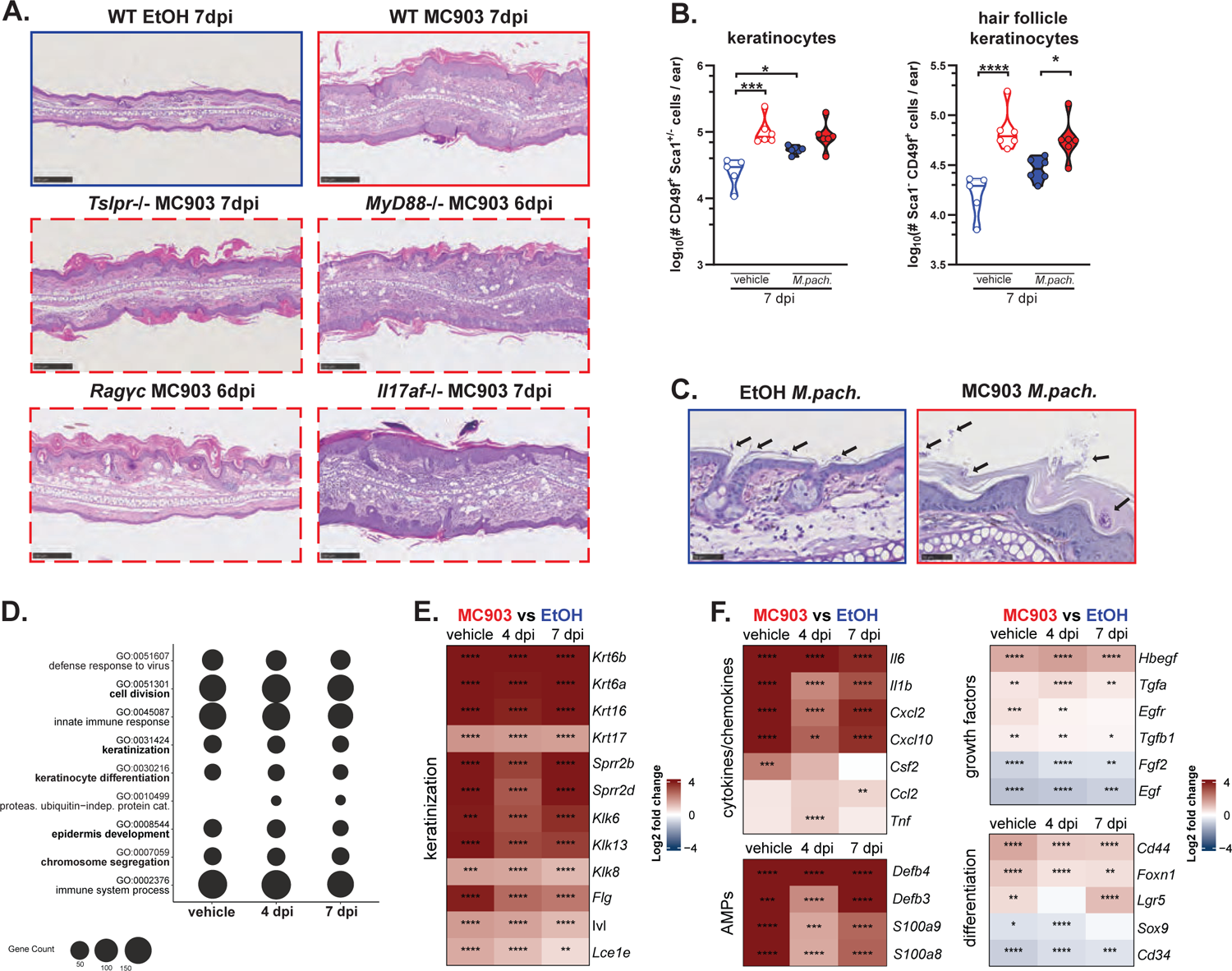
Physical barrier impairment in AD-like skin favors *Malassezia* overgrowth. The ear skin of WT C57BL/6, *Tslpr^-/-^*, *Myd88^-/-^*, *Ragγc^-/-^* and *Il17af^-/-^* mice was treated with MC903 or EtOH and associated with *M. pachydermatis* as in Fig. 1A. **A.** Representative H&E-stained skin sections. **B.** Quantification of overall (left) and hair follicle (right) keratinocytes. **C.** PAS-stained histology sections of WT ear skin with arrows highlighting fungal cell aggregates. **D.-F.** Top upregulated GO terms related to biological process (D) and differentially expressed genes (log2 fold changes, E-F) from the RNA-Seq dataset introduced in Fig. 4D-E comparing MC903-vs. EtOH-treated ear skin that was (4 dpi, 7 dpi) or was not (vehicle) associated with *M. pachydermatis*. Heat maps indicate log2 fold change of selected genes related to keratinization (E) and wound healing (F). Data in B are pooled from two independent experiments with 2-4 mice per group each. Each symbol represents one mouse. The mean of each group is indicated. Statistical significance was determined using two-way ANOVA (B). Stars in heat maps (E, F) reflect statistical significance of FDR values. *p<0.05, **p<0.01, ***p<0.001, ****p<0.0001. **See also Figure S5.**

Scrutinizing our RNA-Seq dataset (Fig. 4E, Fig. S5D) revealed a strong response signature of epidermal barrier impairment and wound healing reminiscent of what has been described for lesional AD.^48^ Keratinization, epithelial cell division, and epidermis development were among the top differentially regulated pathways at all time points (Fig. 5D-F). Importantly, all these MC903-induced changes were independent of the presence of *Malassezia* (Fig. 5D-F). The high significance of the GO term “defense response to virus” appearing at the top of the list (Fig. 5D) is explained by the strong induction of 2’-5’-oligoadenylate synthetases (OAS), which in addition to their antiviral activity also drive cell cycle progression and keratinocyte proliferation in the skin.^49^ Amongst the most highly upregulated genes were genes encoding for structural proteins involved in late keratinocyte differentiation, including involucrin (*Ivl)*, filaggrin (*Flg),* and small proline-rich proteins (*Sprr2b* and *Sprr2d*),^50^ as well as the epithelium-specific serine proteases kallikreins, which are released from injured epithelium to promote desquamation^51^ (Fig. 5E). The expression of inflammatory keratins related to keratinocyte hyperproliferation and stress response (*Krt6a*, *Krt6b*, *Krt16*, and *Krt17*)^52^ was up-regulated as well (Fig. 5E). Dense accumulations of keratohyalin granules containing keratin filaments, filaggrin, and loricrin, highlighted in H&E-stained sections, further illustrated the enhanced keratinization (Fig. S5E). Given the close connection between re-epithelialization and the progression of wound healing, genes related to epidermal repair were differentially regulated in the MC903-treated skin. These include the suprabasal keratinocyte gene *Foxn1*, which encodes a transcription factor regulating the epithelial-mesenchymal transition process,^53^ and the hair follicle stem cell genes *Lgr5*, *Sox9* and *Cd34* ^54^ (Fig. 5F). The epidermal and fibroblast growth factor genes *Egf* and *Fgf2* were downregulated as observed in chronic wounds.^55^ The induction of key genes involved in glycolytic metabolism (Fig. S5F) reflected the high energy demand of proliferating keratinocytes in lesional AD.^56–58^ Many of the inflammatory and antimicrobial genes that we found strongly induced under MC903 conditions are also linked to wound healing.^48,59,60^ This includes small proline-rich proteins mentioned above, which can exhibit antimicrobial function as shown recently.^61^ The strong antimicrobial and inflammatory signature in the AD-like murine skin (Fig. 5F) was surprising considering the observed fungal overgrowth and suggested that *Malassezia* is able to overcome the host defense response by favorable niche conditions that benefit its thriving. In summary, our results reveal little effects of innate and adaptive skin immunity on *Malassezia* overgrowth and of *Malassezia* on epidermal homeostasis but highlights the overarching role of epidermal barrier impairment in fungal overgrowth in AD-like skin.

### Fungal overgrowth is conserved across different models of epidermal barrier disruption

To assess the relevance of epidermal barrier impairment for fungal dysbiosis beyond the MC903 model, we employed a model of genetically-induced epidermal barrier dysregulation owing to the constitutive activation of the antioxidant response factor Nrf2 in keratinocytes.^62^ Of note, *Nfe2l2*, the gene encoding Nrf2, as well as numerous Nrf2 response genes were upregulated in MC903-treated skin (Fig. S5G). Hyperkeratosis and changes in the epidermal lipid envelope in mice expressing a constitutively active Nrf2 mutant under the keratin 5 (K5) promoter (K5-caNrf2+ mice) is accompanied by strong upregulation of *Krt16*, the cornified envelope genes *Slpi* and *Sprr2*, the peroxisome proliferator activator receptor (Ppar) *Ppard* gene, as well as the long-chain fatty acid elongation factor *Elovl4* and *Elovl7* genes.^62^ These transcriptional changes in the skin of K5-caNrf2+ mice parallel the phenotype of MC903-treated skin (Fig. S5F). Four days after skin association with *M. pachydermatis*, K5-caNrf2+ mice showed stable upregulation of *Nrf2* as well as its target gene *Nqo1* (Fig. S6A-B) and increased ear thickness and hyperkeratosis including accumulations of keratohyalin granules in comparison to cre-negative controls (Fig. 6A-C, Fig. S6D). Likewise, the high expression levels of *Sprr2b* and *Krt16* genes in response to the overexpression of Nrf2 and its target genes was conserved during *Malassezia* colonization (Fig. 6B, Fig. S6C). Most importantly, the K5-caNrf2+ mice displayed a significant fungal overgrowth in the ear skin and even more pronounced at the snout, compared to the corresponding control mice (Fig. 6D).

**Fig 6.**
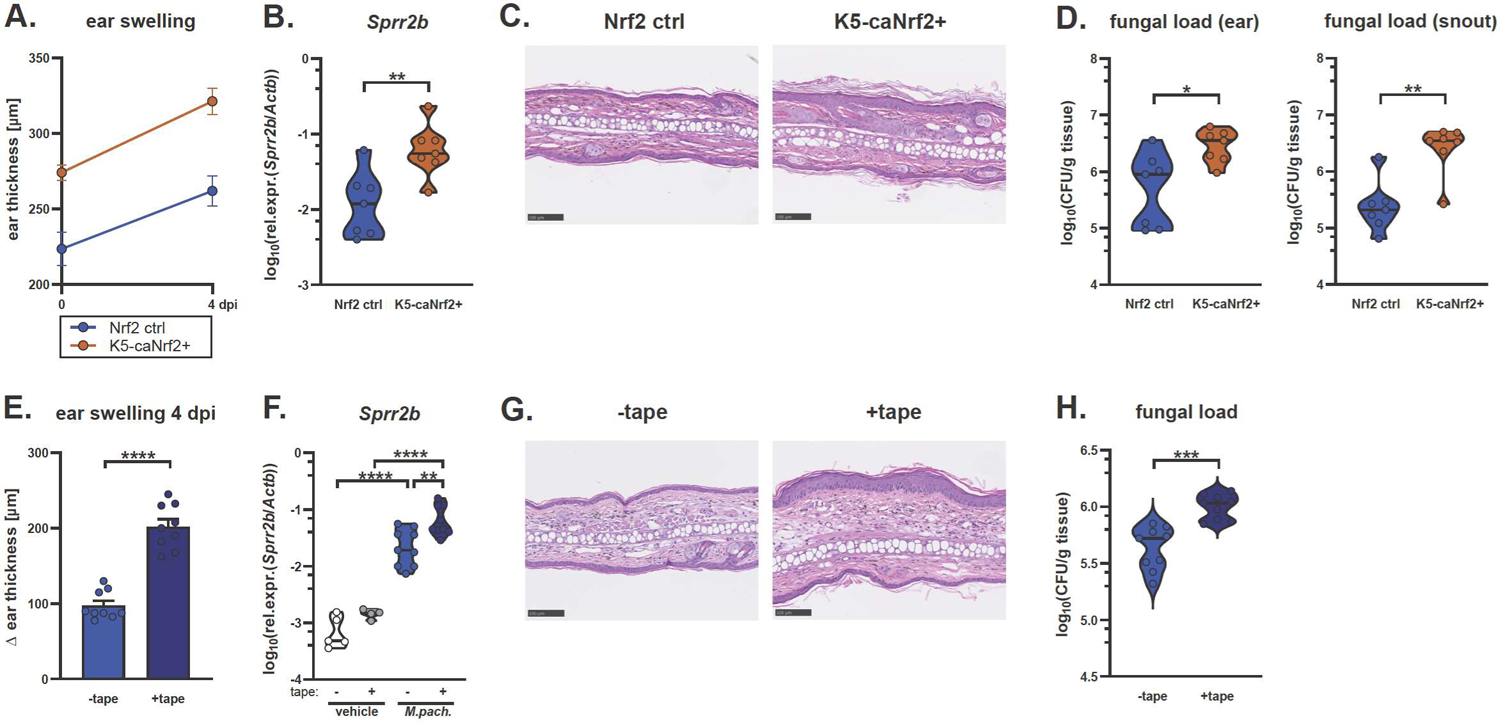
Fungal overgrowth is conserved across different models of epidermal barrier disruption. **A.-D.** The ear skin of K5cre-CMVcaNrf2 (K5-caNrf2+) transgenic and caNrf2 control mice was associated with *M. pachydermatis* for 4 days. Ear thickness (A), *Sprr2b* transcript levels (B), representative H&E-stained skin sections (C), and fungal load in ear skin and snout (D). **E.-H.** The ear skin of WT C57BL/6 mice was tape stripped (+tape) or not (-tape) before association with *M. pachydermatis* for 4 days. Ear thickness (E), transcript levels of *Sprr2b* (F), representative H&E-stained skin sections (G), and skin fungal load (H). Data are pooled from two independent experiments with 2-5 mice per group each. In B, D-F, and H, each symbol represents one mouse, the mean of each group is indicated. The mean +/- SEM per group is indicated in A and E. Statistical significance was determined using Student’s unpaired *t*-test (B, D, E, H) or two-way ANOVA (F). *p<0.05, **p<0.01, ***p<0.001, ****p<0.0001. **See also Figure S6.**

To further assess the relationship of skin barrier impairment with excessive fungal growth, we mechanically disrupted the epidermal barrier of mouse ears by tape stripping prior to association with *Malassezia*. Tape stripping elicits epidermal proliferation and hyperkeratosis as a consequence of the mechanical removal of the stratum corneum,^63,64^ also reflected by increased keratohyalin granule formation (Fig S6E)). In this barrier disrupted skin, introduction of *M. pachydermatis* resulted in strongly exacerbated ear swelling, epidermal hyperplasia, and upregulation of *Sprr2b* and *Krt16* expression (Fig. 6E-G, Fig. S6E-F). Notably, the disruption of the epidermal barrier, once again, caused a clear fungal overgrowth (Fig. 6H). *Malassezia* dysbiosis was independent of elevated *Tslp* expression since fungal skin association led to a downregulation of this cytokine in tape-stripped skin (Fig. S6G). In addition, the observation that *Il17a* expression was not altered in the tape-stripped skin (Fig. S6H) confirms our previous conclusion that fungal overgrowth in AD skin is not primarily a consequence of dysregulated type 17 immunity. Together, our data show that epidermal barrier impairment, regardless of its genetic, chemical, or mechanical origin, strongly favoured *Malassezia* growth and, thereby allowed the fungus to overcome existing antifungal defence mechanisms.

### Changes in lipid metabolism in AD-like skin support *Malassezia* lipid acquisition and growth

Next, we sought to identify factors that benefit *Malassezia* growth in the AD-like skin. Because *Malassezia* relies on extracellular lipids to grow and that AD is characterized by major abnormalities in lipid metabolism, we next studied the relationship between the alterations in lipid metabolism in AD and *Malassezia* overgrowth. The skin of flaky tail (*ft*/*ft*) mice, another widely used model of AD, which is due to naturally acquired mutations in the *Flg* and *Tmem79* genes, manifest an AD-like epidermal barrier defect.^65,66^ We previously detailed the profound changes in the lipid metabolism in these mice, which are characterized by aberrant lamellar body (LB) cargo and secretory system, increased peroxisomal fatty acid oxidation, and accumulation of shorter chain fatty acid and ceramide species at the cost of very-long-chain epidermal lipids.^56^ *ft/ft* mice exposed to *M. pachydermatis* showed enhanced fungal colonization in comparison to WT control mice (Fig. 7A). The difference in epidermal hyperplasia observed at baseline between *ft/ft* and WT control mice was conserved upon *Malassezia* association (Fig 7B-C), albeit overall skin thickness increased after fungal association as seen before in other AD-like models (Fig. 6A, E). Likewise, the hyperkeratosis-related genes *Sprr2b* and *Krt16* were not differentially expressed in *Malassezia*-associated *ft*/*ft* and WT mice (Fig. S7A-B), while dysregulation in cutaneous lipid metabolism and hypoxia, representing hallmarks of the AD-like phenotype in *ft*/*ft* mice,^56^ were conserved (Fig. 7D, Fig. S7C). The differential regulation of the master regulators of lipid metabolism, namely enhanced expression of *Ppard* and decreased expression of *Ppara* and *Pparg*, as well as the transcriptional changes in fatty acid shuttling (*Fabp5*, *Crot*), uptake (*Slc27a2*), and synthesis/elongation (*Elovl1*, *Elov4*, *Elov6*) genes in the skin of *ft*/*ft* mice^56^ paralleled transcriptional changes in MC903-treated mice as well as in Nrf2-overexpressing mice (Fig. 7E, S7D-E). Notably, these changes in lipid metabolism were independent of the type 2 phenotype in the MC903 model (Fig S7D). Altogether, our results suggest that *Malassezia* takes advantage of the altered lipid metabolism in AD skin, which is affecting the epidermal lipid composition and barrier integrity, thereby, providing the fungus with a more readily accessible niche for its thriving.

**Fig 7.**
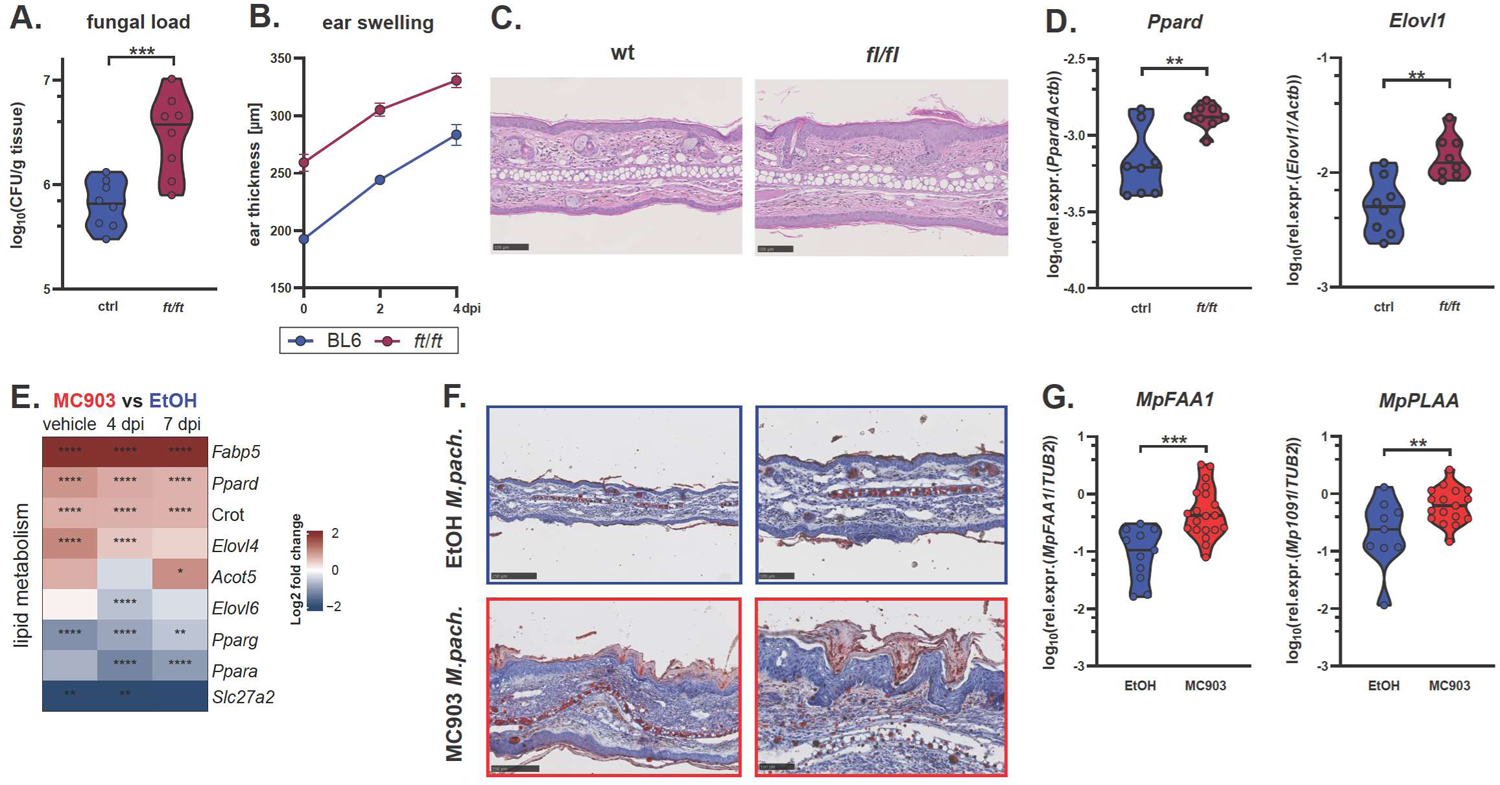
Changes in lipid metabolism in AD-like skin support *Malassezia* lipid acquisition and growth. **A.-D.** Association of the ear skin of flaky tail (*ft*/*ft*) and WT C57Bl/6 control mice with *M. pachydermatis* for four days. Skin fungal load (A), ear thickness kinetics (B), representative H&E-stained skin sections (C), and skin expression levels of the lipid metabolism genes *Ppard* and *Elovl1* (D). **E.** Heat maps indicating selected differentially expressed genes (log2 fold changes) related to lipid metabolism from the RNA-Seq dataset introduced in Fig. 4D-E comparing MC903-vs. EtOH-treated ear skin. **F.-G.** The ear skin of WT C57BL/6 mice was treated with MC903 or EtOH and associated with *M. pachydermatis* or vehicle for 4 days as in Fig. 1A. Oil Red staining at two different magnifications (F) and fungal gene expression in the skin (G). Data are pooled from two (A-D) or four (G) independent experiments with 2-5 mice per group each. Each symbol in A, D and H represents one mouse, lines indicate the mean of each group. The mean +/- SEM is indicated in B. Statistical significance was determined using Student’s unpaired *t*-test (A, D, H). Stars in heat map (E) reflect statistical significance of FDR values. *p<0.05, **p<0.01, ***p<0.001, ****p<0.0001. **See also Figure S7.**

*Malassezia* relies on lipid acquisition from the host since these lipid-dependent yeasts lack fatty acid synthase for de novo lipid synthesis.^67^ Lipids were increased at the site of *Malassezia* colonization in the stratum corneum including hair follicles (Fig. 5C), as revealed by Oil Red staining of the MC903-treated skin (Fig. 7F). Importantly, we found enhanced metabolic activity of the fungus in AD-like skin that was manifested by altered gene expression compared to control skin. Notably, *MpFAA1*, a key fungal gene implicated in fatty acid uptake, activation and consequently in fungal growth,^67,68^ was upregulated in the AD-like skin (Fig. 7H). We further found increased expression of the fungal phospholipase A2 activating gene *MpPLAA* (Fig. 7G). Phospholipases are involved in the acquisition of fatty acids from the host environment.^69^ The significance of this result is supported by the finding that *M. pachydermatis* isolates from dogs with dermatitis or otitis display enhanced production and activity of phospholipase A2, among other phospholipases.^70–72^ These transcriptional changes in *Malassezia* suggest increased lipid uptake in the AD-like skin providing the fungus with a growth advantage under conditions of increased nutrient availability. Together, these findings suggest that the fungus benefits from the altered skin environment.

## DISCUSSION

Microbial dysbiosis and functional adaptations in the colonizing microbes is a common phenomenon in diverse pathological skin conditions.^2,73^ Although the causal relationship between dysbiosis and pathogenesis remains largely unclear in most cases, a better understanding of what drives dysbiosis may provide opportunities for restoring homeostasis of protective barrier tissues and improving health. Alterations in colonization and function of *Malassezia*, the by far most abundant fungal commensal yeast on our skin, are associated with a range of skin conditions including dandruff, pityriasis versicolor, seborrheic dermatitis, and atopic dermatitis.^14^

Here we show that a dysfunctional cutaneous barrier predisposes for unrestrained growth of *Malassezia*. While homeostatic immunity is a central pillar of immunosurveillance of commensal organisms, changes in the immune status in AD-like skin such as overt type 2 and reduced type 17 cytokine production were insufficient to explain *Malassezia* dysbiosis. Instead, we found that AD-related dysregulation of the epidermal barrier provides a favourable environment for fungal thriving with enhanced accessibility and changes in lipid composition of the metabolic niche allowing for excessive *Malassezia* growth. These findings provide mechanistic insights into conditions underlying *Malassezia* dysbiosis in several common skin pathologies characterized by partially overlapping skin barrier defects, including AD and ichthyosis. They further advance the understanding of factors ensuring fungal commensalism in the healthy skin.

Skin disorders have a high global prevalence. Although usually not lethal, they significantly limit the well-being of those affected and consequently are listed the 4^th^ greatest non-fatal disabling condition, with fungal skin infections, acne, pruritus, and AD ranking among the most frequent skin pathologies.^74,75^ New immunotherapies have become available for some of these conditions including AD that alleviate the burden significantly in those responding. However, they are not effective in all patient subgroups and there is a need for effective patient stratification and understanding of disease promoting factors.^76^ As such, secondary bacterial and fungal infections are a complicating factor in skin barrier diseases, which can actively contribute to pathogenesis of disease as has been studied extensively in case of *S. aureus* colonizing AD lesional skin.^11^

In this study, we observed excessive *Malassezia* growth in four different experimental models of skin barrier dysregulation. Barrier disruption is a defining feature of AD and related diseases such as ichthyosis vulgaris, which share a similar decrease of *FLG* expression.^77^ Associated alterations in the lipid organisation of the stratum corneum and the lamellar body secretory pathway in AD patients have been described in humans and dogs.^78–80^ These aspects of altered barrier lipid metabolism are closely recapitulated in the *ft*/*ft* and MC903 models of AD.^56^ The changes include a shift of epidermal lipids towards shorter chain fatty acids and ceramides (C_16_-C_22_), at the expense of those harbouring C_24_ and C_26_.^81–83^ The enriched lipids are within the spectrum of fatty acids that are preferentially utilised by *Malassezia* (C_12_-C_20_).^84,85^ Due to the lack of fatty acid synthetases, *Malassezia* depends on the assimilation of fatty acids from the host by means of lipases, phospholipases, and sphingomyelinases.

Our results indicate that *Malassezia* adapts to the altered niche conditions in AD-like skin and uses them to its own advantage. *M. pachydermatis* responds to altered lipid availability in barrier dysregulated skin by increased expression of genes involved in enhanced lipid acquisition, such as *MpFAA1* and *MpPLAA*. Long-chain fatty acyl-CoA synthetase 1 (FAA1) recognizes and facilitates uptake of C_14_-C_16_ fatty acids important for *Malassezia* growth.^67,68^ PLAA in turn positively regulates cytosolic and calcium-independent phospholipase A2 activities. Phospholipases are a heterogenous group of enzymes that hydrolyse ester linkages in phospholipids by cleaving specific ester bonds.^86^ Importantly, PLA2 activity was found increased in *M. pachydermatis* isolates from skin lesions of dogs presenting dermatitis and/or otitis, as reported by three independent studies.^70–72^ Phospholipases are considered virulence factors in many pathogenic fungi due to their potential to damage host cell membranes,^86^ which can further aggravate barrier disruption and cause inflammation by inducing host cell lysis and subsequent release of danger-associated molecular patterns. In addition, the hydrolysis of lipids by lipases may affect the acidic pH of the skin, as it is the case for bacterial lipases,^87,88^ while enzyme activity itself is affected by pH.^89,90^ Therefore, enhanced phospholipase expression by *Malassezia* may allow the fungus to better thrive on the host skin but may also contribute to disease pathogenesis. Another known fungal determinant associated with skin inflammation is secreted aspartyl protease 1 (SAP1), which was found to be highly expressed in AD lesional skin.^91^ The pro-inflammatory role of SAP1 in *Malassezia* pathogenicity was confirmed by means of a gene deficient mutant in *M. furfur* that was impaired in driving inflammation in barrier disrupted skin.^91^

That a dysregulation of the skin barrier establishes a favorable skin environment for *Malassezia* is supported by *Malassezia* overgrowth in canine AD^92–94^ and hereditary ichthyosis^95^. In further support of the causal relationship between loss of barrier integrity and enhanced fungal control, a case report found topical treatment improving the skin barrier function to coincide with a reduction in the fungal load.^96^ No consistent data exist for differential *Malassezia* colonization levels on human skin.^13,97^ Amplicon and metagenomic sequencing studies generally report relative abundances of individual genera or species. Meanwhile, older culture-based studies suffer from the difficulty of cultivating *Malassezia* spp. and from small cohort sizes. Finally, the method of sampling has not been standardized regarding the method, sampling site and intensity of swabbing or tape stripping.

Besides the defining feature of skin barrier dysregulation, AD most often manifests as an allergic disease.^98^ In sensitized AD patients, anti-*Malassezia* IgE levels correlate with AD severity^99,100^ and *Malassezia*-reactive Th2 cells are enriched, although their role in AD pathogenesis remains uncertain.^23,30^ In MC903-treated experimental animals, we have not detected an enrichment of *Malassezia*-responsive Th2 cells upon fungal association, possibly due to the protocol of a single administration of a relatively high fungal load. Experimental induction of eczema by house dust mite or of fungal airway allergy relies on repeated administration of a low dose inoculum.^101,102^ Of note, type 2 immunity was not responsible for the observed fungal overgrowth based on results obtained with MC903-treated TSLPR-deficient mice and from IL-4cx injection experiments. Importantly, the impaired barrier function and altered lipid profile in MC903-treated skin was also independent of TSLP signalling. Likewise, in the MC903 model, acute itch was shown to rely on infiltrating neutrophils, which are again induced independently of TSLP signalling in response to MC903.^103^ Together, this might indicate a discrepancy between the experimental model of AD in mice and the human disease where the type 2 immune axis appears to play a more prominent role in the disease. Namely, in human AD patients, targeting the IL-4/13 receptor by Dupilumab is effective in 60-65% of patients by alleviating pruritus with a positive effect on barrier integrity as a result of reduced scratching among other factors.^104,105^

Besides type 2 immunity, immune dysregulation in the atopic skin in the MC903 model is hallmarked by reduced antifungal type 17 response. Although our results manifest a partial contribution of the compromised IL-17 production in the MC903-treated skin, the pronounced fungal overgrowth persisted irrespective of the IL-17 status of the host. In addition, expression of antimicrobial peptides, many of which are known to be regulated by IL-17, are not suppressed in MC903-treated skin but rather increased, in line with the overall strong inflammatory phenotype. As such, *Malassezia* can overcome enhanced antimicrobial defense and inflammation and effectively access favorable nutrients in the altered niche that foster enhanced colonization. As such, our results identify an important role of MC903 in the modulation of the skin microbiome/mycobiome, which should be considered for pharmacological use of calcipotriol and other Vitamin D analogues used against skin carcinomas^106,107^ and psoriasis (Dovonex®)^44,45^. Therapeutic approaches to strengthen the epidermal barrier function can support the restoration of a homeostatic interaction with commensal microorganisms and prevent overt growth and aberrant expression of virulence factors. Maintaining stable commensalism is vital for host physiology given that fungal commensals influence many aspects of host health, barrier protection, host defense, and even social behavior.^108^

## STAR METHODS

### Key Resource Table

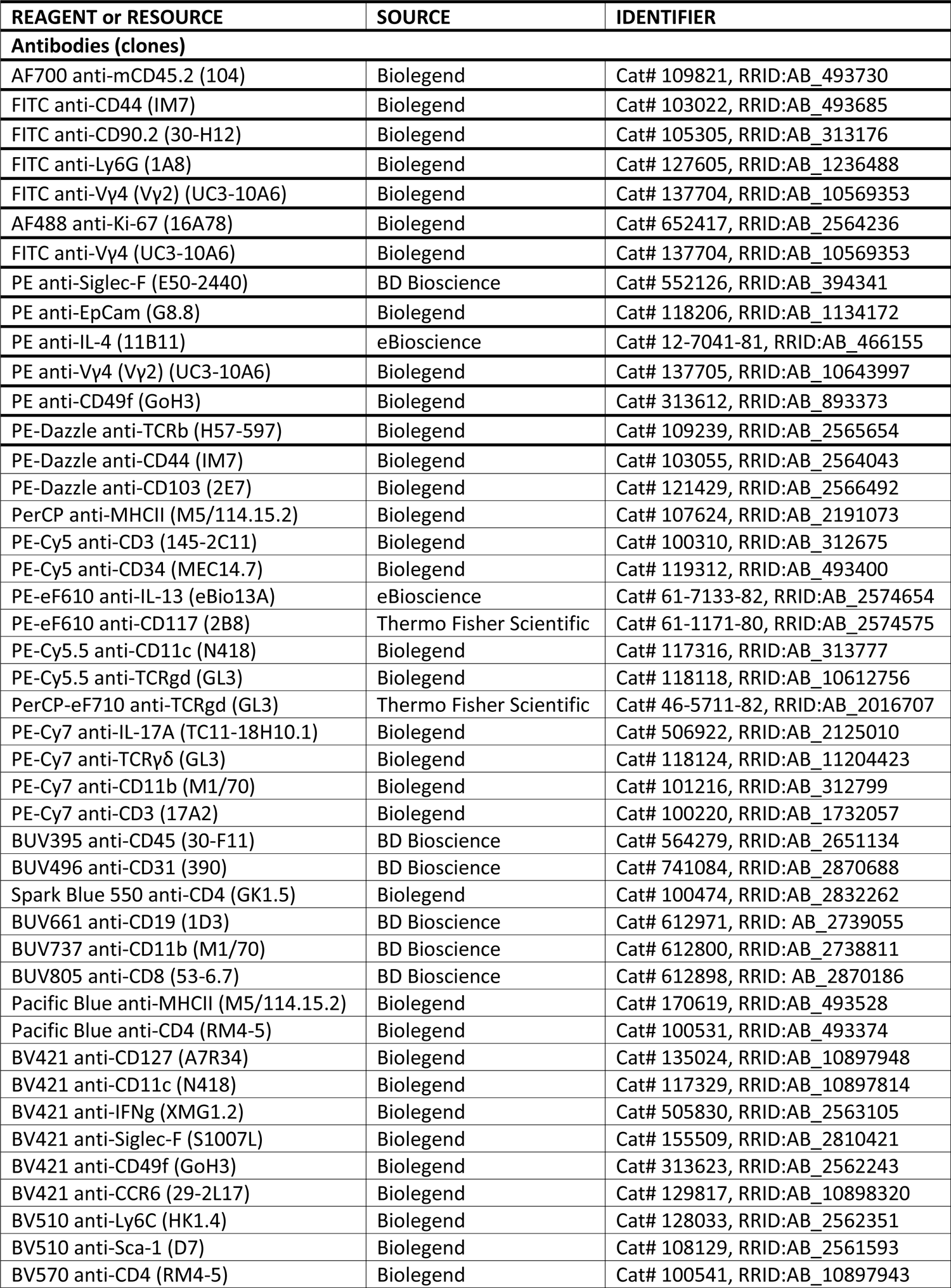
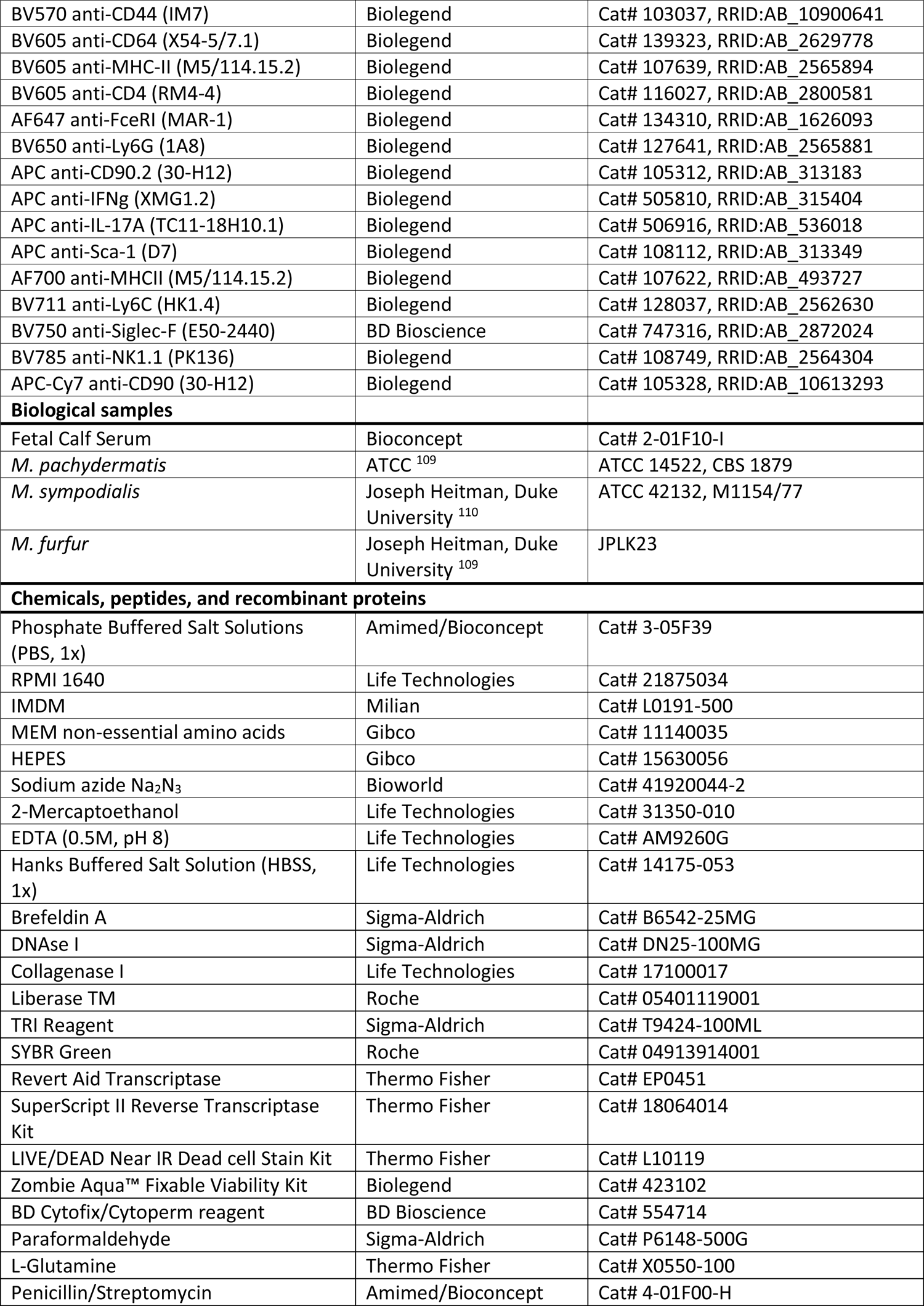
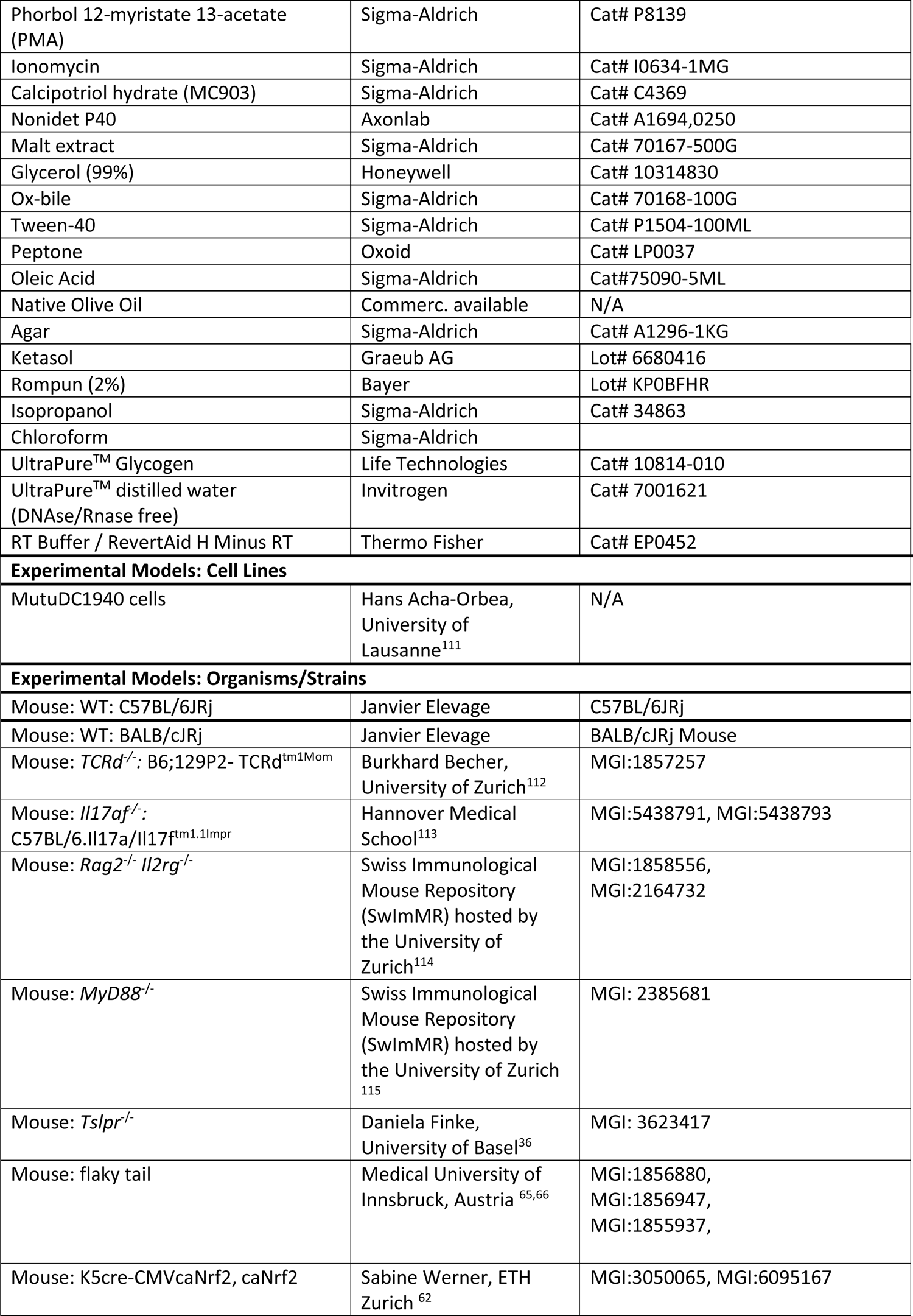
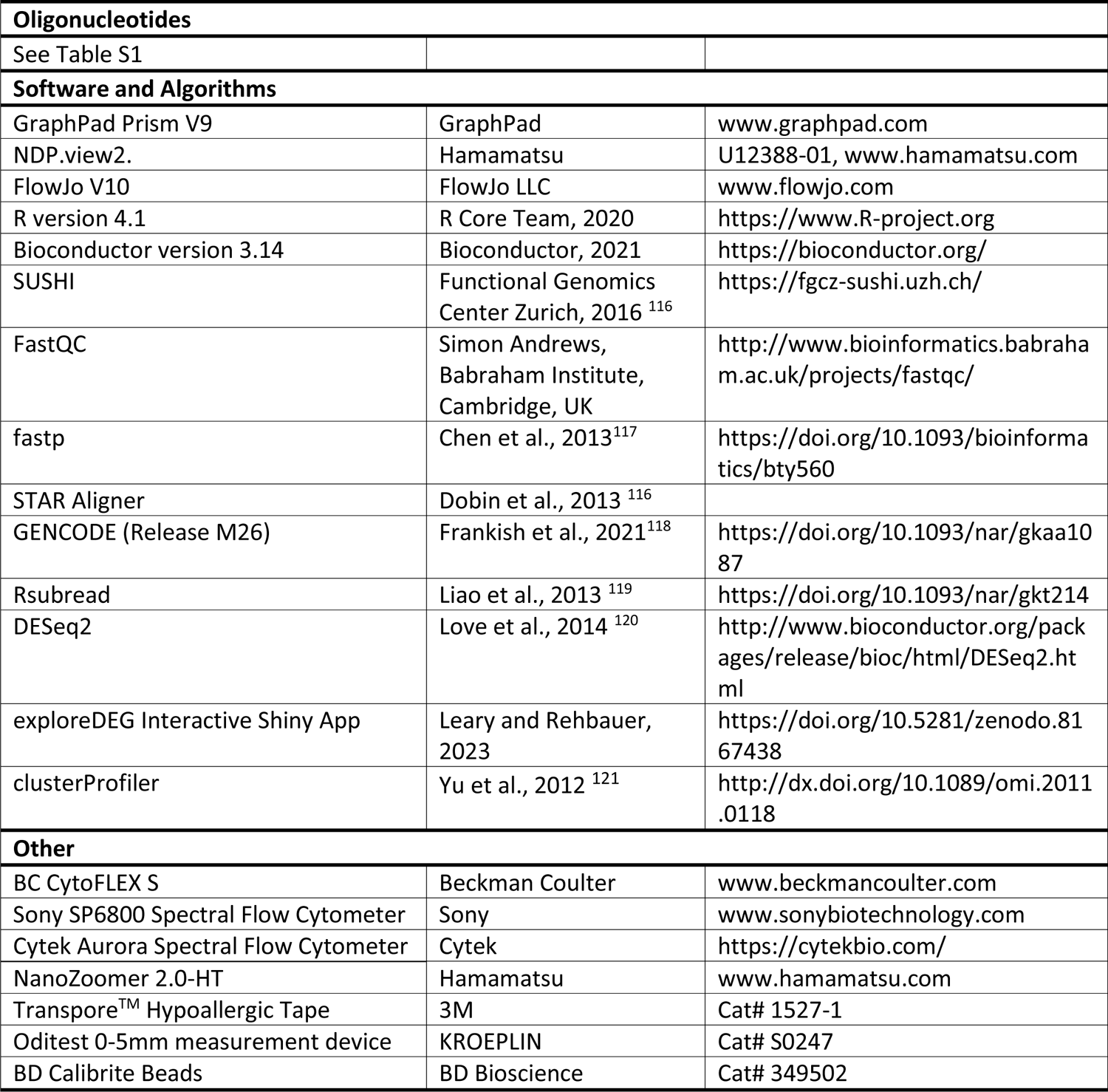

#### Contact for Reagent and Resource Sharing

Further information and requests for resources and reagents should be directed to and will be fulfilled by the lead contact Salomé LeibundGut-Landmann. *Tslpr*^-/-^ mice were obtained via an MTA between University of Zurich and the NHLBI/NIH. The MutuDC1940 cell line was obtained via an MTA between the University of Zurich and the University of Lausanne.

### Experimental Model and Subject Details

#### Ethics statement

All mouse experiments in this study were conducted in strict accordance with the guidelines of the Swiss Animals Protection Law and were performed under the protocols approved by the Veterinary office of the Canton Zurich, Switzerland (license number 168/2018 and 142/2021). All efforts were made to minimize suffering and ensure the highest ethical and humane standards according to the 3R principles.^122^

#### Animals

WT C57Bl/6JRj and Balb/c mice were purchased from Janvier Elevage (France). *Il17af^-/-^* ^113^, *Tcrd^-/-^* ^112^, *Rag2*^-/-^ *Il2rg*^-/-^ ^114^ (kindly obtained from the Swiss Immunological Mouse Repository), and *MyD88*^-/-^ mice ^115^ were bread at the Laboratory Animal Science Center of the University of Zurich (LASC). *Il17af^-/-^*mice were used with littermate and co-housed WT control mice (*Il17af^wt^*). *MyD88*^-/-^ mice were used with heterozygous littermate controls. Flaky tail (*ft/ft*) mice ^65,66^ were bred at the Medical University of Innsbruck, Austria, and used for experiment with WT controls from the same facility. *Tslpr*^-/-^ mice^36^ (kindly obtained from Daniela Finke) were bred at the Department of Biomedicine of the University of Basel and used for experiment with WT controls from the same facility. K5cre-CMVcaNrf2 and caNrf2 control mice ^62^ (kindly obtained from Sabine Werner) were bred at the ETH Zurich Phenomics Center. All mice were on the C57BL/6 background and kept under specific pathogen-free conditions. All experiments were conducted at the Laboratory Animal Science Center of the University of Zurich. Female as well as male mice were used at 8-15 weeks of age in sex- and age-matched groups. Groups of vehicle-treated and infected animals were kept in separate cages to avoid cross-contamination.

#### Fungal strains

All *Malassezia* strains used in this study were used previously in the Leibundgut lab.^30^ *M. pachydermatis* strain ATCC 14522 (CBS 1879)^123^ was originally purchased from ATCC. *M. sympodialis* strain ATCC 42132 ^110^ and *M. furfur* strain JPLK23 (CBS 14141)^123^ were originally obtained from Joseph Heitman (Duke University Medical Center, Durham, NC). All strains were grown in liquid modified Dixon (mDixon) medium (Malt Extract (Sigma Aldrich), desiccated Ox-bile (Sigma Aldrich), Tween-40 (Sigma Aldrich), Peptone (Oxoid), Glycerol (Honeywell), Oleic Acid (Sigma Aldrich)) at 30°C and 180 rpm for 2-3 days. Heat-killed *Malassezia* was prepared by twice washing Malassezia grown from liquid culture in PBS and exposing to 85°C for 45 min at a concentration of OD5_A600_/ml.

### Experimental models

#### MC903-induced model of AD

Induction of an AD-like skin phenotype by MC903 was based on Moosbrugger-Martinz et al.^31^ In short, both ears of mice were treated with MC903 (calcipotriol hydrate; Sigma) in ethanol or with pure ethanol as a control (Sigma) on the dorsal and ventral side of both ears. The dose of MC903 was 1.125 nmol in 25µl ethanol per ear. The treatment was done daily for five days and again four days after a resting period of two days. The treatment was continued after infection on the ventral ear side only on day 2 (experiments with endpoint at 4 dpi) or daily from day 2 to 5 (experiments with endpoint at 7 dpi). For experiments with Balb/c mice, left ears were treated with EtOH while the right ear was treated with MC903. Ear thickness was continuously monitored using the Oditest S0247 0-5 mm measurement device (Kroeplin).

#### Systemic treatment of mice with IL-4cx

IL-4cx, composed of 0.5 µg mouse IL-4 (PeproTech) complexed with 5 µg anti–IL-4 mAb (clone 11B11; Bio X cell) was injected i.p. daily from day−2 until day 0 as described previously.^124^ Mice received further IL-4cx injections on days 2, 4, and 6 post-infection with *Malassezia*.

#### Tape stripping of mouse ear skin

Tape stripping was performed by repeated application and removal of adhesive tape (Transpore™ Hypoallergenic, 3M; 10 times per ear). Ear thickness was measured using the Oditest S0247 0-5 mm measurement device (Kroeplin).

#### Epicutaneous association of mouse ear skin with *Malassezia*

Epicutaneous infection of the mouse ear skin was performed as described previously.^30,33^ *Malassezia* strains grown in mDixon medium were washed in PBS and suspended in native olive oil at a density of 10 OD_A600_/ml. 100 µL suspension (corresponding to 1 OD_A600_ or 5×10^6^ CFU/ml) of yeast cells was applied topically onto the dorsal ear skin while mice were anaesthetized. For determining the fungal loads in the skin, tissue was transferred in water supplemented with 0.05 % Nonidet P40 (AxonLab), homogenized with a TissueLyzer (Qiagen) for 6 minutes at 25 Hz, plated on mDixon agar, and incubated at 30°C for 3 to 4 days.

## Method details

### Histology

Mouse tissue was fixed in 4% PBS-buffered paraformaldehyde overnight and embedded in paraffin. Sagittal sections (2-8µm) were stained either with hematoxylin and eosin (H&E) or with Periodic-acidic Schiff (PAS) reagent and counterstained with Hematoxylin and mounted with Pertex (Biosystem, Switzerland) according to standard protocols. Oil red staining was performed and counterstained with Hemalum solution and mounted on aqueous mounting medium according to standard protocols. All images were acquired with a digital slide scanner (NanoZoomer 2.0-HT, Hamamatsu) and analyzed with NDP view2.

### RNA isolation and RT-qPCR

Isolation of total RNA from murine ear skin was performed using TRI reagent (Sigma-Aldrich) according to the manufacturer’s protocol with the modification that the tissue was homogenized using a TissueLyzer (Qiagen) for 6 minutes at 25 Hz before extraction of the nucleic acids with chloroform. For isolation of fungal RNA, an additional step of homogenization with glass beads (0.5 mm, Sigma) was included twice for 2 minutes at 30 Hz interrupted by 30 s cooling periods on ice. cDNA was generated by RevertAid reverse transcriptase (ThermoFisher) and random nonamers. Quantitative PCR was performed using SYBR green (Roche) and a QuantStudio 7 Flex (Life Technologies) or a 7500 Real-Time PCR System (Applied Biosystems) instrument. The primers used for qPCR are listed in Table S1. All RT-qPCR assays were performed in duplicate, and the relative expression (rel. expr.) of each gene was determined after normalization to mouse *Actb* or *M. pachydermatis TUB2* housekeeping genes. Fungal cDNA data was excluded when expression of the housekeeping gene was above a Ct value of 40 or when unspecific amplification occurred.

### Bulk RNA-Seqencing

RNA sequencing was performed by the Functional Genomics Center Zürich. RNA was extracted using the Direct-zol RNA MiniPrep Plus Kit (Qiagen) following manufacturer’s protocol. Extracted RNA was prepared for sequencing using the Illumina TruSeq RNA assay following manufacturer’s protocol. Sequencing was performed on the Illumina NovaSeq 6000 using the S1 Reagent Kit v1.5 (100 cycles) as per manufacturer’s protocol. Demultiplexing was performed using the Illumina bcl2fastq Conversion Software. Individual library size ranged from 20.6 million to 28.5 million reads.

### RNA-Seq data analysis

RNA sequencing analysis was performed using the SUSHI framework,^116^ which encompassed the following steps: Read quality was inspected using FastQC, and sequencing adaptors removed using fastp; Alignment of the RNA-Seq reads using the STAR aligner^125^ and with the GENCODE mouse genome build GRCm39 (Release M26) as the reference^118^; the counting of gene-level expression values using the ‘featureCounts’ function of the R package Rsubread^119^; differential expression using the generalized linear model as implemented by the DESeq2 Bioconductor R package^120^, and; Gene Ontology (GO) term pathway analysis using the hypergeometric over-representation test via the ‘enrichGO’ function of the clusterProfiler Bioconductor R package^121^. Plots were created using the SUSHI framework including the exploreDEG Interactive Shiny App. All R functions were executed on R version 4.1 (R Core Team, 2020) and Bioconductor version 3.14. Data will be made publicly available upon acceptance of the manuscript for publication.

### Isolation of murine skin and lymph node cells

For digestion of total ear skin, mice ears were removed, cut into small pieces, and transferred into Hank’s medium (Ca^2+^- and Mg^2+^-free, Life Technology), supplemented with Liberase TM (0.15 mg/mL, Roche) and Dnase I (0.12 mg/mL, Sigma-Aldrich) and incubated for 1 hour at 37°C. Ear-draining lymph nodes (dLN) were digested with DNAse I (2.4mg/ml Sigma-Aldrich) and Collagenase I (2.4mg/ml, Roche) in PBS for 15 min at 37°C. Both cell suspensions were filtered through a 70 μm cell strainer (Falcon) and rinsed with PBS supplemented with 5 mM EDTA (Life Technologies), 1% fetal calf serum and 0.02% NaN_3_. Single cell suspensions were used for re-stimulation or stained directly for flow cytometry.

### *Ex vivo* T cell stimulation

For *in vitro* re-stimulation of skin T cells, skin cell suspensions were incubated in a U-bottom 96-well plate (1/4 ear per well) with cRPMI medium (RPMI supplemented with 10% fetal calf serum, HEPES, sodium pyruvate, non-essential amino acids, β-mercaptoethanol, Penicillin and Streptomycin) with phorbol 12-myristate 13-acetate (PMA, 50 ng/ml, Sigma-Aldrich) and ionomycin (500 ng/ml, Sigma-Aldrich) for 5 h at 37 °C, 5% CO_2_ in the presence of Brefeldin A (10 μg/ml). For *in vitro* re-stimulation of lymph node T cells, 10^6^ lymph node cells per well of a flat bottom 96-well plate were co-cultured with 1×10^5^ DC1940 cells ^111^ that were pulsed with 2.5×10^5^ heat-killed fungal cells. Brefeldin A (10 mg/ml, Sigma-Aldrich) was added for the last 4 hours of incubation at 37°C, 5% CO_2_.

### Flow cytometry

Single cell suspensions were stained with antibodies as listed in the Key Resource Table. LIVE/DEAD Near IR stain (Life Technologies) or Zombie Aqua fixable viability staining (Biolegend) was used for exclusion of dead cells. After surface staining, cells were fixed and permeabilized with Cytofix/Cytoperm (BD Biosciences) for subsequent intracellular staining in Perm/Wash buffer (BD Bioscience). All staining steps were carried out on ice. Data were acquired on a Spectral Analyzer SP6800 (Sony), CytoFLEX S (Beckman Coulter) or 5L Cytek Aurora (Cytek). All fcs files were analyzed with FlowJo software V10 (FlowJo LLC). Gating of the flow cytometric data was performed according to the guidelines for the use of flow cytometry and cell sorting in immunological studies,^126^ including pre-gating on viable and single cells for analysis. Absolute cell numbers were calculated based on a defined number of counting beads (BD Bioscience, Calibrite Beads) that were added to the samples before flow cytometric acquisition.

### ELISA

Blood was collected by post-mortem cardiac puncture and transferred to blood collection tubes (BD Microtainer^®^, BD) for serum isolation. Detection of IgE from mouse serum was performed by ELISA using ELISA MAX™ Standard Set Mouse IgE (Biolegend) according to the manufacturer’s instructions.

### Quantification and Statistical Analysis

Statistical significance was determined by unpaired Student’s *t*-test with Welch’s correction and one- or two-way ANOVA with Dunnet’s or Tukey’s multiple comparison test, as appropriate, using GraphPad Prism software. Data displayed on a logarithmic scale were log-transformed before statistical analysis.Significance is indicated as follows: *p< 0.05; **p<0.01; ***p<0.001; ****p<0.0001.

## DATA AVAILABILITY

All raw data and metadata linked to this study will be made publicly available on Zenodo (https://zenodo.org/) and the relevant doi number will be indicated here upon acceptance of the manuscript for publication.

## ACKNOWLEDGMENT

The authors would like to thank Daniela Finke (University of Basel, Department of Biomedicine, Basel, Switzerland) for *Tslpr*^-/-^ mice; Sabine Werner and Michael Koch (Institute for Molecular Health Sciences, ETH Zurich, Switzerland) for K5cre-CMVcaNrf2 mice; Onur Boyman for anti-IL-4 antibody; the staff of the Laboratory Animal Service Center of University of Zurich for animal husbandry; staff of the Laboratory for Animal Model Pathology of University of Zurich for histology; Peter Leary from the Functional Genomics Center Zürich for bioinformatics support; members of the LeibundGut-lab and of SKINTEGRITY.CH for helpful advice and discussions. This work was supported by the Swiss National Science Foundation (grant # 310030_189255, to S.L.-L.) and by a Candoc grant from the University of Zürich (grant #5843, to F.R.).

## AUTHOR CONTRIBUTIONS

F.R. and S.L.-L. designed the study and wrote the original draft of the manuscript. F.R. performed the experiments and analysed the data. P.Z. developed the multiparameter flow cytometry panel in Fig. 2 and S2 and analysed the data together with F.R.. S.L.-L. oversaw all study design and data analysis and acquired funding. S.D. provided flaky tail mice and important intellectual input; All authors discussed the results, and reviewed and edited the manuscript.

## SUPPLEMENTARY FIGURE LEGENDS

**Fig S1.** (related to Figure 1). Atopic skin environment leads to *Malassezia* overgrowth after epicutaneous association in the MC903 model. **A.-C.** The ear skin of WT C57BL/6 mice was treated with MC903 or EtOH and associated with *M. pachydermatis* or vehicle as in Fig. 1A. The dorsal (dors.) and ventral (vent.) side of the ear were analyzed separately for expression of *Cyp24a1*, *Tslp*, and *Il17a* transcripts (A), fungal load (B), or immune cell numbers (C) at 7 dpi. **D.** MC903 (red) or EtOH (blue)-treated ear skin of WT C57BL/6 mice was associated with *M. furfur* or *M. sympodialis* (*M.symp.*) and skin fungal load (CFU) was determined at 6 or 4 dpi, respectively. **E.-G.** The ear skin of WT Balb/c mice was treated with MC903 or EtOH and associated with *M.pach.* or vehicle, as in Fig. 1A. Ear thickness kinetics (E), *Tslp* transcript levels (F), and skin fungal load (G) at 2 dpi. Data in A-C and E-F are from one experiment with 2-3 mice per group. Data in D are from one representative of two independent experiments with 2-4 mice per group each. In A-D and F-G, each symbol represents one mouse, the mean of each group is indicated. In E, each symbol is the mean +/- SD per group and time point. Statistical significance was determined using two-way ANOVA (A, B, C left), one-way ANOVA (C, F), or Student’s unpaired *t*-test (D, G). ns p>0.1, *p<0.05, **p<0.01.

**Fig S2.** (related to Figure 2). MC903-induced skin inflammation is characterized by broad infiltration of hematopoietic cell subsets. The ear skin of WT C57BL/6 mice were treated with MC903 or EtOH and associated with *M. pachydermatis* or vehicle as in Fig. 1A. **A.** Gating strategy for cell subset identification. A representative example of MC903-treated skin at 7 dpi is shown. **B.** Quantification of the indicated cell populations as in Fig. 1C. Each symbol represents one mouse. The mean of each group is indicated. Data are pooled from two independent experiments with three mice per group each. Statistical significance was determined using two-way ANOVA. *p<0.05, **p<0.01, ***p<0.001, ****p<0.0001.

**Fig S3.** (related to Figure 3). Fungal overgrowth in the MC903-treated skin is independent of type 2 immunity. **A-E.** The ear skin of WT C57BL/6 mice was treated with MC903 or EtOH and associated with *M. pachydermatis* as in Fig. 1A. Cytokine production by skin and draining lymph node (dLN) T cells was assessed at 7 dpi after *ex vivo* re-stimulation. Representative FACS plots (A) showing IL-4, IL-13, and IL-17A-production by CD44^+^ CD4^+^ T cells after re-stimulation with either dendritic cells pulsed with heat-killed *M. pachydermatis* (DC+*hkM.pach.*) or with PMA and ionomycin as indicated. Percentages of events in each quadrant are indicated. Quantification of IL-4^+^ (B) and IL-13^+^ (C) CD44^+^ CD4^+^ dLN T cells or of IL-4^+^ CD44^+^ CD4^+^ skin T cells (D). Absolute numbers and percentages are shown **E.** Quantification of CD45^+^ cells in the MC903 or EtOH-treated and *Malassezia*-associated ear skin of *Tslpr*-/- and WT control mice at 7 dpi. **F.** Representative H&E-stained skin sections of mice treated with IL-4cx or PBS as in Fig. 3B at the indicated time point after *M. pachydermatis* association or vehicle treatment. In B-D data are from one (C, D), one of two (E) or two (B) independent experiments with 3-4 mice per group each. Each symbol represents one mouse, the mean per group is indicated. Statistical significance was determined using two-way ANOVA (B, D, E) or Student’s unpaired *t*-test (C). ns p>0.1, *p<0.05, **p<0.01, ****p<0.0001.

Fig S4. (related to Figure 4). Fig 4. Decreased type 17 immunity in the MC903-exposed skin alone is not responsible for fungal overgrowth. **A.** Gating strategy for Vγ4^+^ γδ T cells and CD4^+^ T cells in the ear skin after *ex vivo* restimulation with PMA and ionomycin. **B.** Quantification of IFN-γ^+^ CD90^+^ CD44^+^ cell numbers in the MC903- or EtOH-treated and *M. pachydermatis* (*M.pach.*)-associated ear skin of WT C57BL/6 mice after *ex vivo* re-stimulation with PMA and ionomycin at 6 dpi. **C.** Quantification of IL-17A^+^ CD90^+^ CD44^+^ cells (left), IL-17A^+^ CD44^+^ CD4^+^ T cell (middle), and IL-17A^+^ CD44^+^ Vγ4^+^ γδ T cell subsets (right) in the lymph nodes of MC903- or EtOH-treated and *Malassezia*-associated WT C57BL/6 mice after *ex vivo* re-stimulation with dendritic cells pulsed with heat-killed *M. pachydermatis* at 7 dpi. **D.-E.** *Il17a* transcript levels in the MC903 or EtOH-treated and *Malassezia*-associated ear skin of *Tslpr*^-/-^ and WT control mice at 7 dpi (D) or of *MyD88*^-/-^ and *MyD88*^+/-^ control mice at 6 dpi (E). **F.** *Il17a* transcript levels in the ear skin of WT C57BL/6 mice that were injected with IL-4cx and associated with *M. sympodialis* (*M.symp.*) as in Fig. 3B at 2 and 7 dpi. DL, detection limit. Data are from one representative of two independent experiments (B, D), from a single experiment (E), or pooled from two independent experiments (F) with 2-5 mice per group each. Each symbol corresponds to one mouse, the mean of each group is indicated. Statistical significance was determined using Student’s unpaired *t*-test (B-C) or two-way ANOVA (D-F). *p<0.05, **p<0.01, ***p<0.001, ****p<0.0001.

**Fig S5.** (related to Figure 5). Physical barrier impairment in AD-like skin favors *Malassezia* overgrowth. **A.-B.** Gating strategy for keratinocyte subsets (A) and quantification of the indicated subsets in the MC903 or EtOH-treated and *Malassezia*-associated ear skin of WT C57BL/6 mice. **C.** median fluorescence intensity (MFI) of Ki-67 (left) and CD44 (right) expression in overall keratinocytes in the *Malassezia*-associated ear skin of WT C57Bl/6 mice. **D.** Differentially expressed genes from the RNAseq dataset introduced in Fig. 4D-E comparing MC903-vs. EtOH-treated skin at the indicated time points after fungal association (dpi) or vehicle administration. Venn diagram (D) of differentially up- and downregulated genes (Log2 fold change>2, FDR<0.05). **E.** H&E-stained skin sections of *M. pachydermatis*-associated WT control and *Tslpr*^-/-^ mice treated with MC903 or EtOH as in Fig. 3E. **F.**-**G.** Heat map from the RNAseq dataset introduced in Fig. 4D-E indicating selected differentially expressed genes (log2 fold changes) related to glycolysis (F) and Nrf2 regulation (G). Data are from one (C) or two (B) independent experiments with 2-5 mice per group each. Each symbol represents one mouse. The mean +/- SD of each group is indicated in C. Statistical significance was determined using Student’s unpaired *t*-test (C) or two-way ANOVA (B). Stars in heat maps (F, G) reflect statistical significance of FDR value. *p<0.05, **p<0.01, ***p<0.001, ****p<0.0001.

**Fig S6.** (related to Figure 6) Fungal overgrowth is conserved across different models of epidermal barrier disruption. **A.-C.** *Nfe2l2* (A), *Nqo1* (B), and *Krt16* (C) transcript levels in the ear skin of K5cre-caNrf2 (K5-caNrf2+) transgenic and caNrf2 control mice that were associated with *M. pachydermatis* for four days. **D.-E.** H&E-stained skin sections of *M. pachydermatis*-associated K5-caNrf2+ mice (D) and of tape stripped WT C57BL/6 mice 4 days after *M. pachydermatis* association. **F.-H.** *Krt16* (F)*, Tslp* (G), and *Il17a* (H) transcript levels in the ear skin of WT C57BL/6 mice that were tape stripped (+tape) or not (-tape) before association with *M. pachydermatis* for 4 days. Data are from two independent experiments with 2-5 mice per group each. Each symbol represents one mouse. Statistical significance was determined using Student’s unpaired *t*-test (A-C) or two-way ANOVA (F-H). *p<0.05, ****p<0.0001.

**Fig S7.** (related to Figure 7) Changes in lipid metabolism in AD-like skin support *Malassezia* lipid acquisition and growth. **A.-C.** *Sprr2b* (A), *Krt16* (B), and *Hif1a* (C) transcript levels in the *M. pachydermatis*-associated ear skin of flaky tail (*ft*/*ft*) and WT control mice at 4 dpi. **D.-E.** Expression levels of genes associated with epidermal lipid metabolism in the ear skin of *Tslpr*^-/-^ and WT control mice treated with MC903 as in Fig 3E (D) and of K5-caNrf2+ and Nrf2 control mice (E). Data are from one of two independent (D) or pooled from two independent experiments (A-C, E) with 3-5 mice per group each. Each symbol represents one mouse, and the mean of each group is indicated. Statistical significance was determined using two-way ANOVA (B) or Student’s unpaired *t*-test (C-F). *p<0.05, **p<0.01, ***p<0.001, ****p<0.0001.

