## Supplementary figures and images for "Epidermal barrier dysregulation in atopic skin predisposes for excessive growth of the allergy-associated yeast *Malassezia*"

### Supplementary Figures

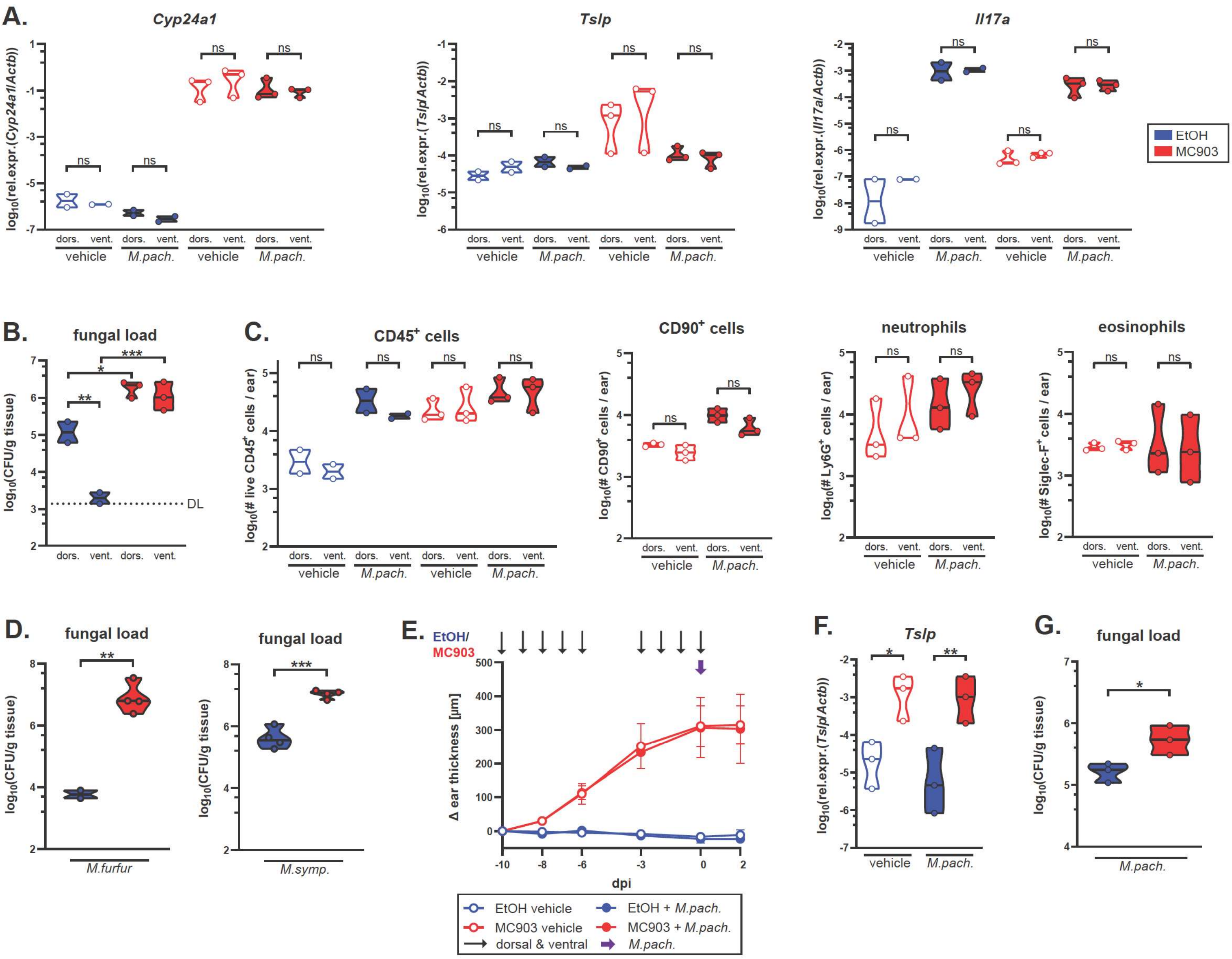

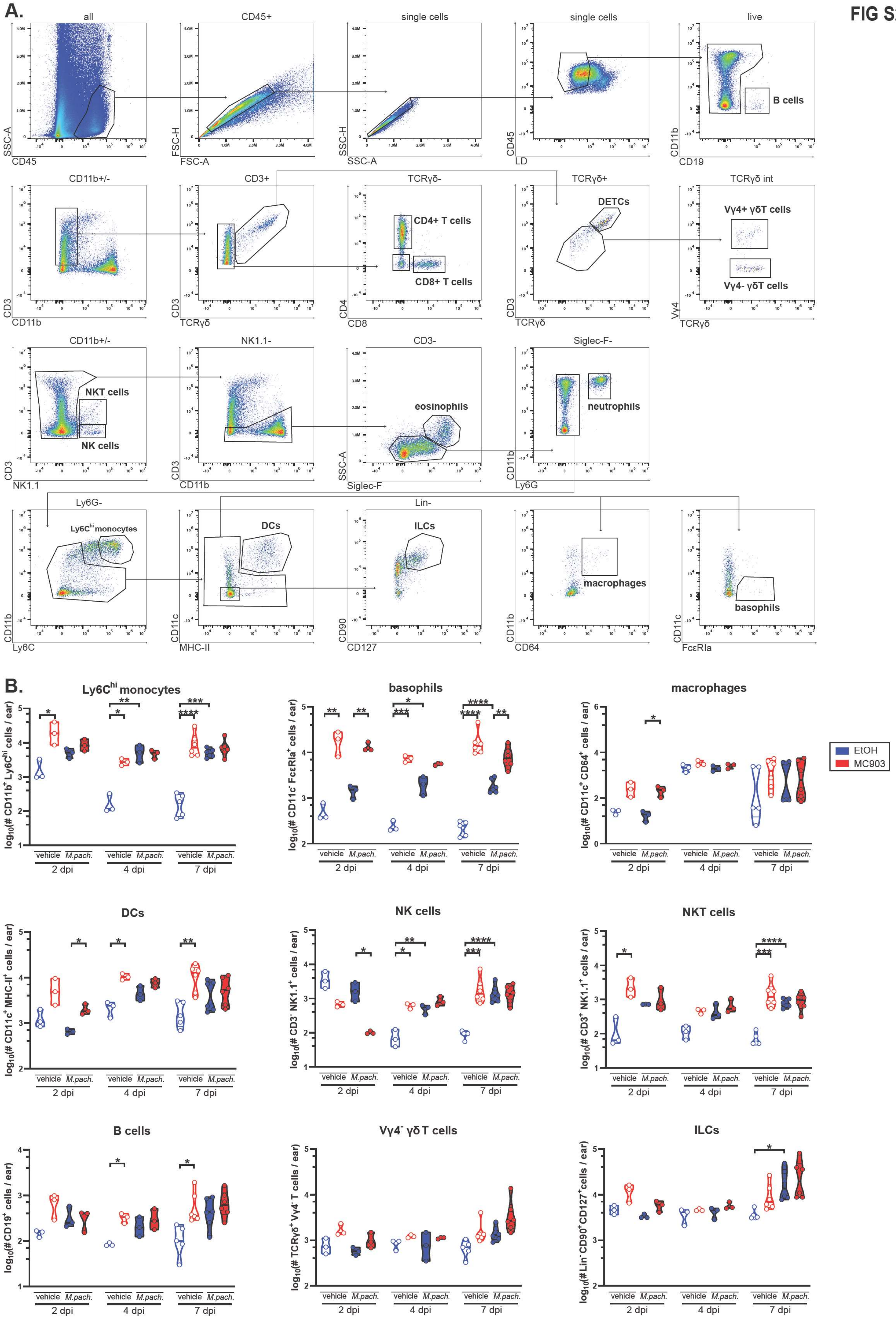

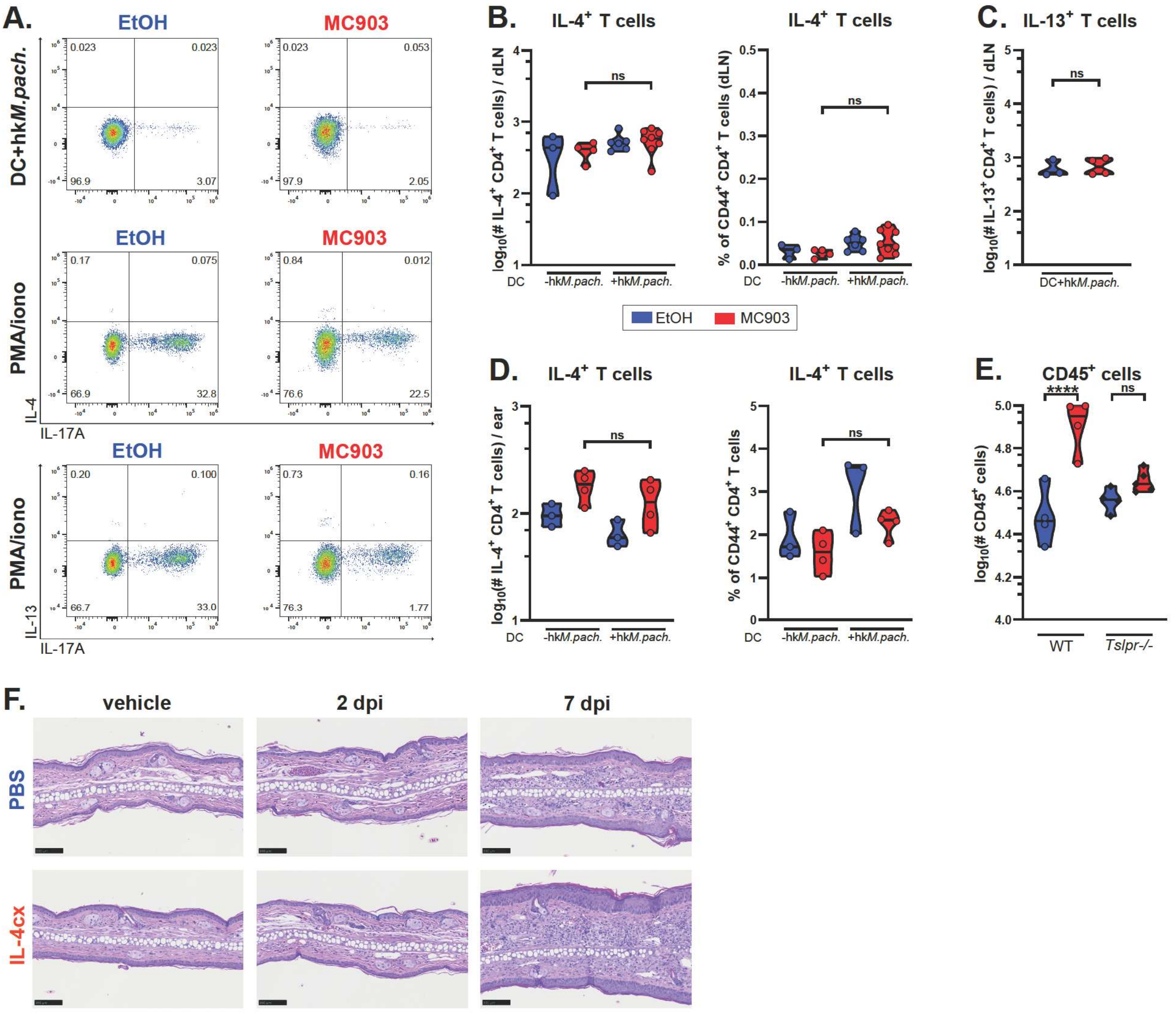

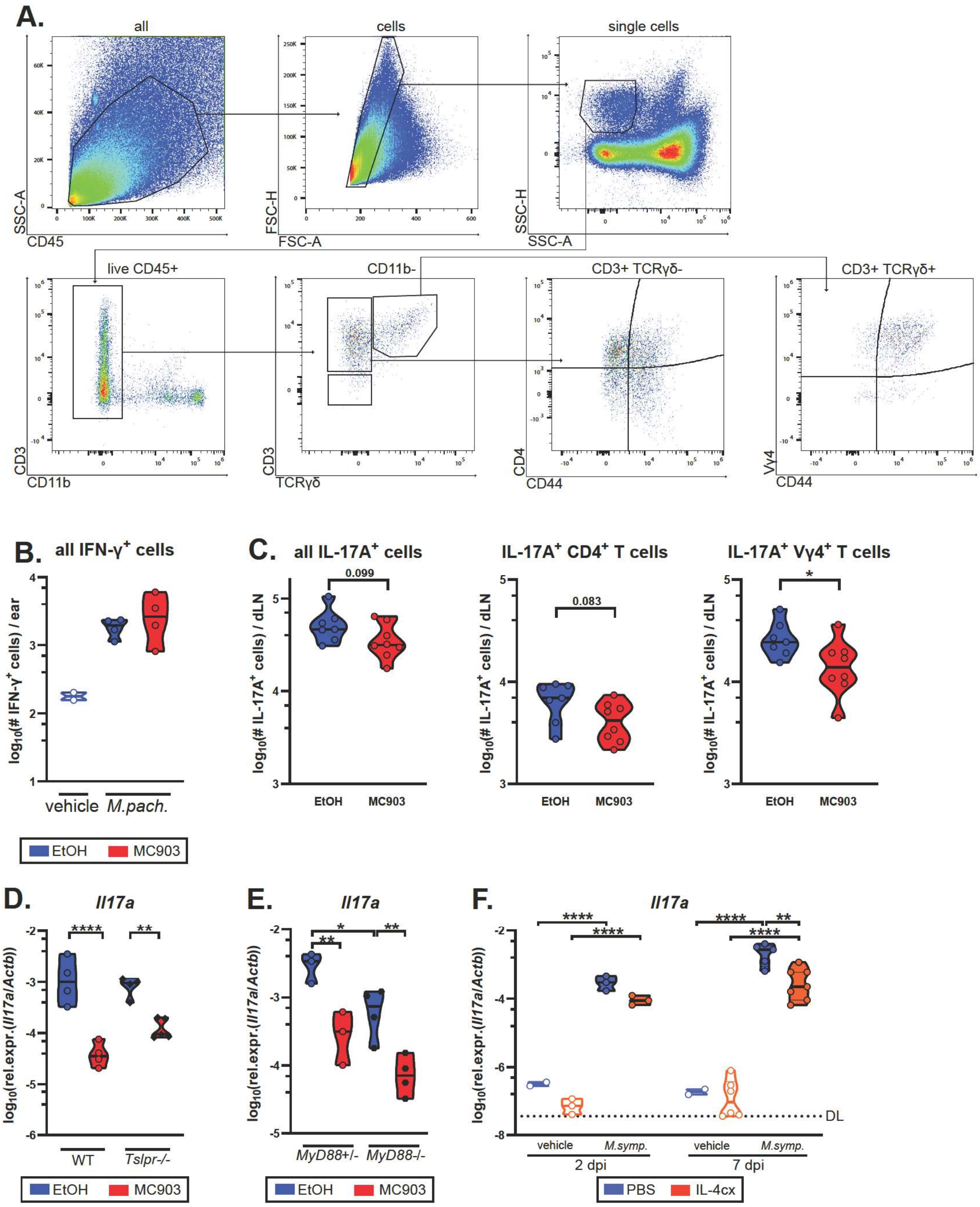

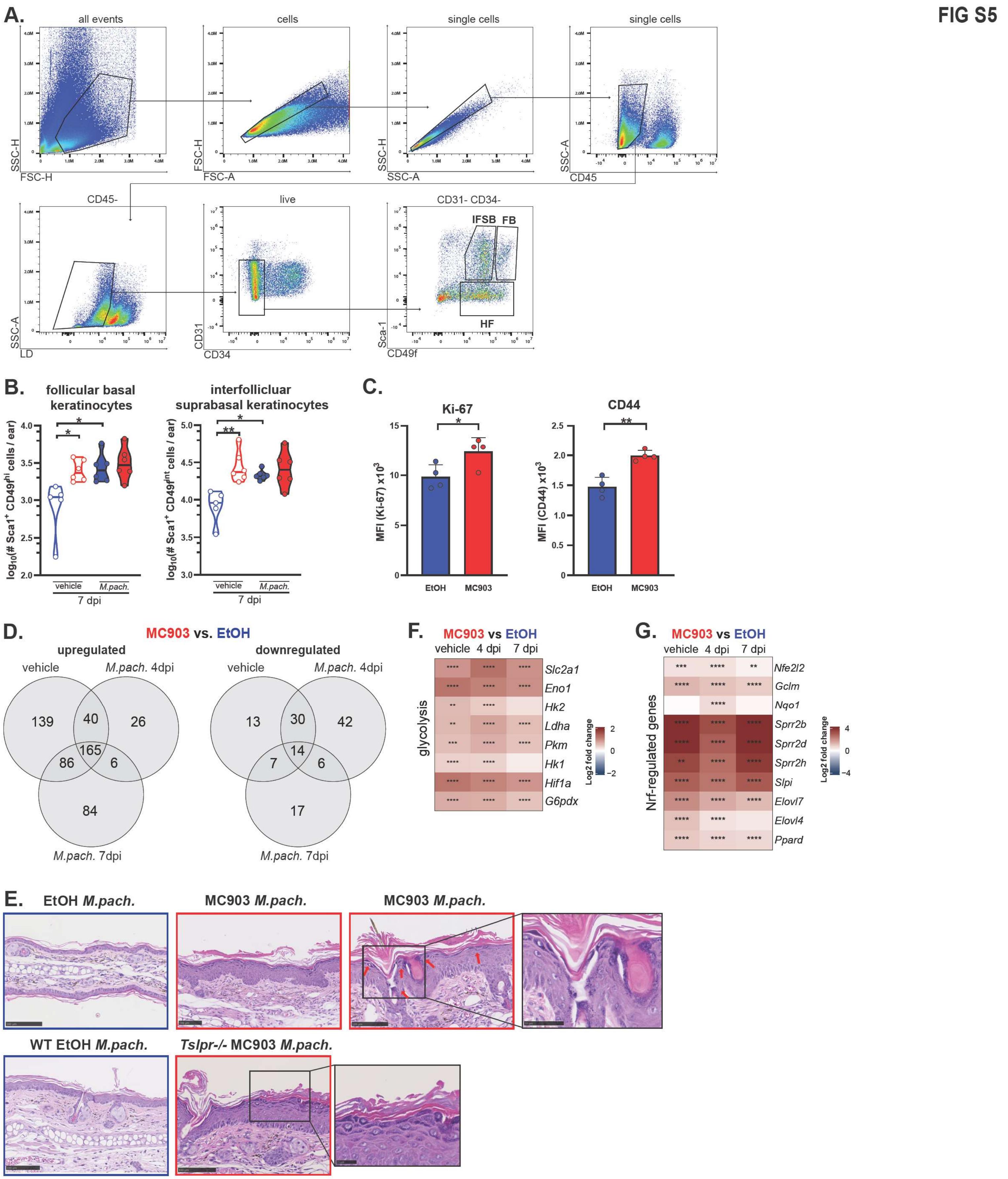

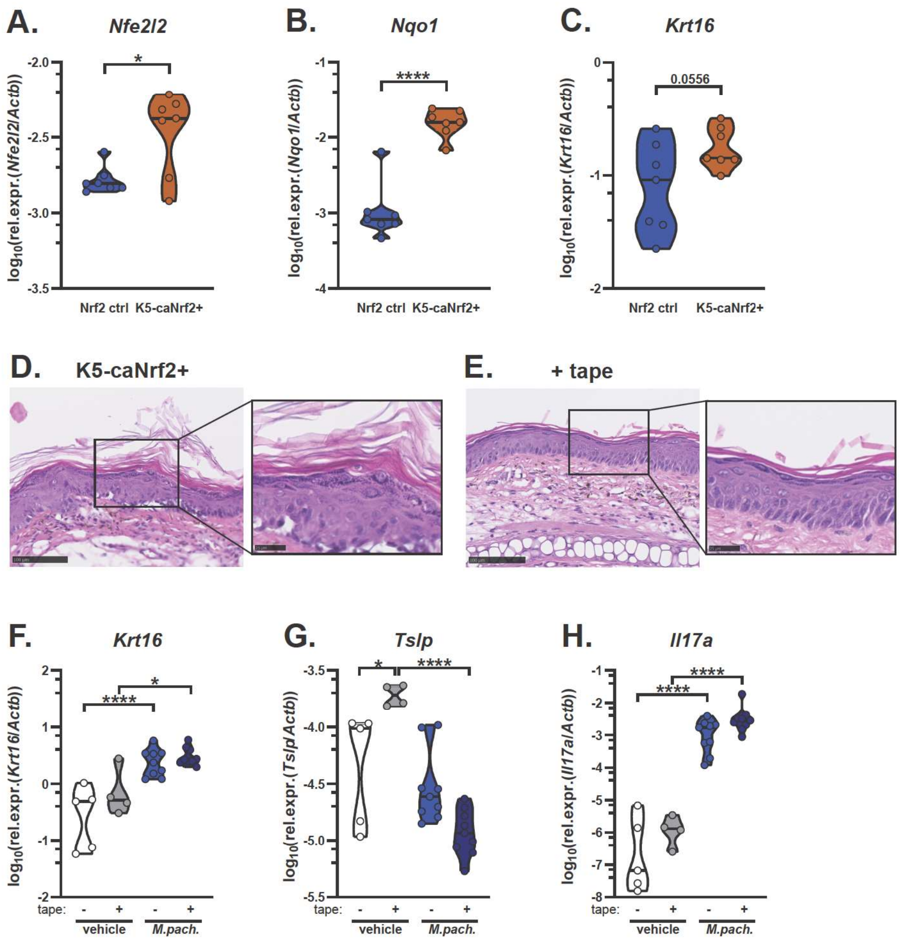

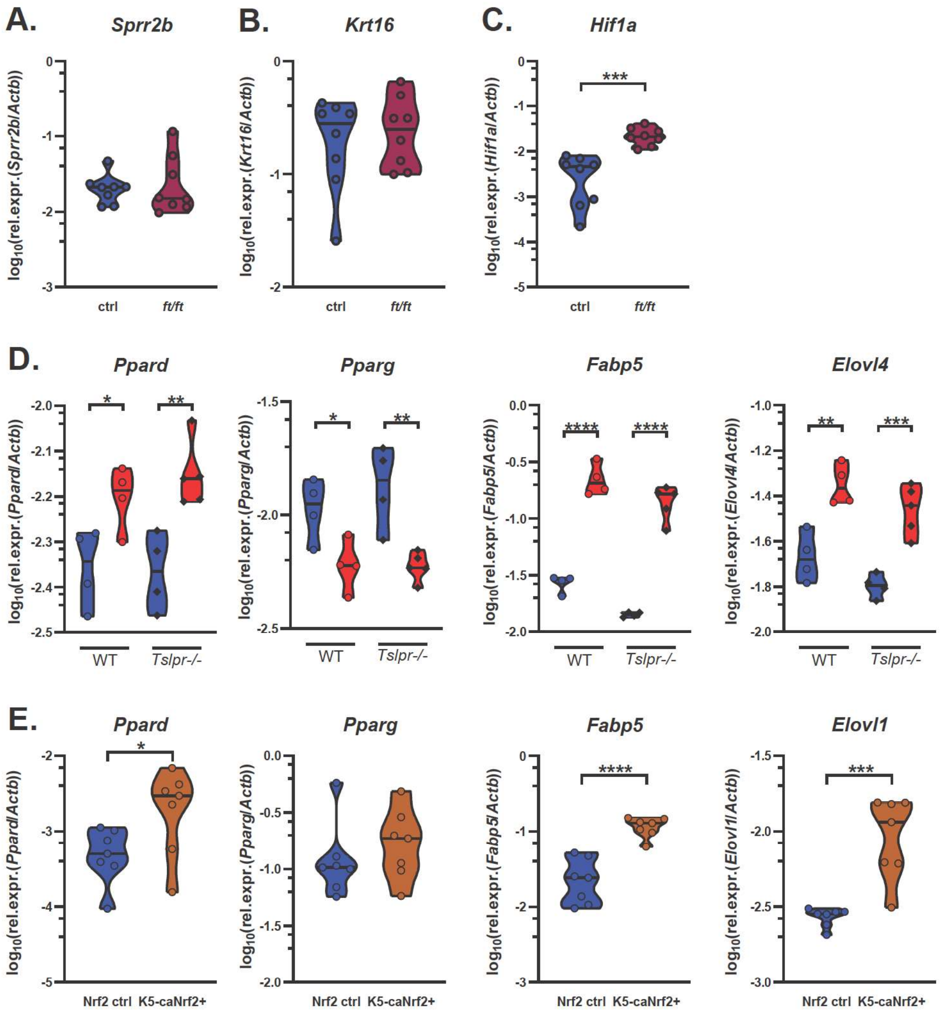
